# scNym: Semi-supervised adversarial neural networks for single cell classification

**DOI:** 10.1101/2020.06.04.132324

**Authors:** Jacob C. Kimmel, David R. Kelley

**Affiliations:** Calico Life Sciences, LLC, South San Francisco, CA, 94080

**Keywords:** single cell, neural network, cell type classification, semi-supervised learning, adversarial learning

## Abstract

Annotating cell identities is a common bottleneck in the analysis of single cell genomics experiments. Here, we present scNym, a semi-supervised, adversarial neural network that learns to transfer cell identity annotations from one experiment to another. scNym takes advantage of information in both labeled datasets and new, unlabeled datasets to learn rich representations of cell identity that enable effective annotation transfer. We show that scNym effectively transfers annotations across experiments despite biological and technical differences, achieving performance superior to existing methods. We also show that scNym models can synthesize information from multiple training and target datasets to improve performance. In addition to high performance, we show that scNym models are well-calibrated and interpretable with saliency methods.

## Introduction

Single cell genomics allows for simultaneous molecular profiling of thousands of diverse cells and has advanced our understanding of development [1], aging [2, 3, 4], and disease [5]. To derive biological insight from these data, each single cell molecular profile must be annotated with a cell identity, such as a cell type or state label. Traditionally, this task has been performed manually by domain expert biologists. Manual annotation is time consuming, somewhat subjective, and error prone. Annotations influence the results of nearly all downstream analysis, motivating more robust algorithmic approaches for cell type annotation.

Automated classification tools have been proposed to transfer annotations across data sets [6, 7, 8, 9, 10, 11, 12]. These existing tools learn relationships between cell identity and molecular features from a training set with existing labels without considering the unlabeled target dataset in the learning process. However, results from the field of semi-supervised representation learning suggest that incorporating information from the target data during training can improve the performance of prediction models [13, 14, 15, 16]. This approach is especially beneficial when there are systematic differences – a domain shift –between the training and target datasets. Domain shifts are introduced between single cell genomics experiments when cells are profiled in different experimental conditions or using different sequencing technologies.

Here, we introduce a cell type classification model that uses semi-supervised and adversarial machine learning techniques to take advantage of both labeled and unlabeled single cell datasets. We demonstrate that this model offers superior performance to existing methods and effectively transfers annotations across different animal ages, perturbation conditions, and sequencing technologies. Additionally, we show that our model learns biologically interpretable representations and offers well-calibrated metrics of annotation confidence that can be used to make new cell type discoveries.

## Results

### scNym

In the typical supervised learning framework, the model touches the target unlabeled dataset to predict labels only after training has concluded. By contrast, our semi-supervised learning framework trains the model parameters on both the labeled and unlabeled data in order to leverage the structure in the target dataset, whose measurements may have been influenced by myriad sources of biological and technical bias and batch effects. While our model uses observed cell profiles from the unlabeled target dataset, at no point does the model access ground truth labels for the target data. Ground truth labels on the target dataset are used exclusively to evaluate model performance.

scNym uses the unlabeled target data through a combination of MixMatch semi-supervision [16] and by training a domain adversary [17] in an iterative learning process (Fig. 1A, Methods). The MixMatch semi-supervision approach combines MixUp data augmentations [18, 19] with pseudolabeling of the target data [20, 15] to improve generalization across the training and target domains. At each training iteration, we “pseudolabel” unlabeled cells using predictions from the classification model, then augment each cell profile using a biased weighted average of gene expression and labels with another randomly chosen cell (Fig. 1B). We fit the model parameters to minimize cell type classification error on these mixed profiles, encourag-ing the model to learn a general representation that allows for interpolation between observed cell states.

**Figure 1:**
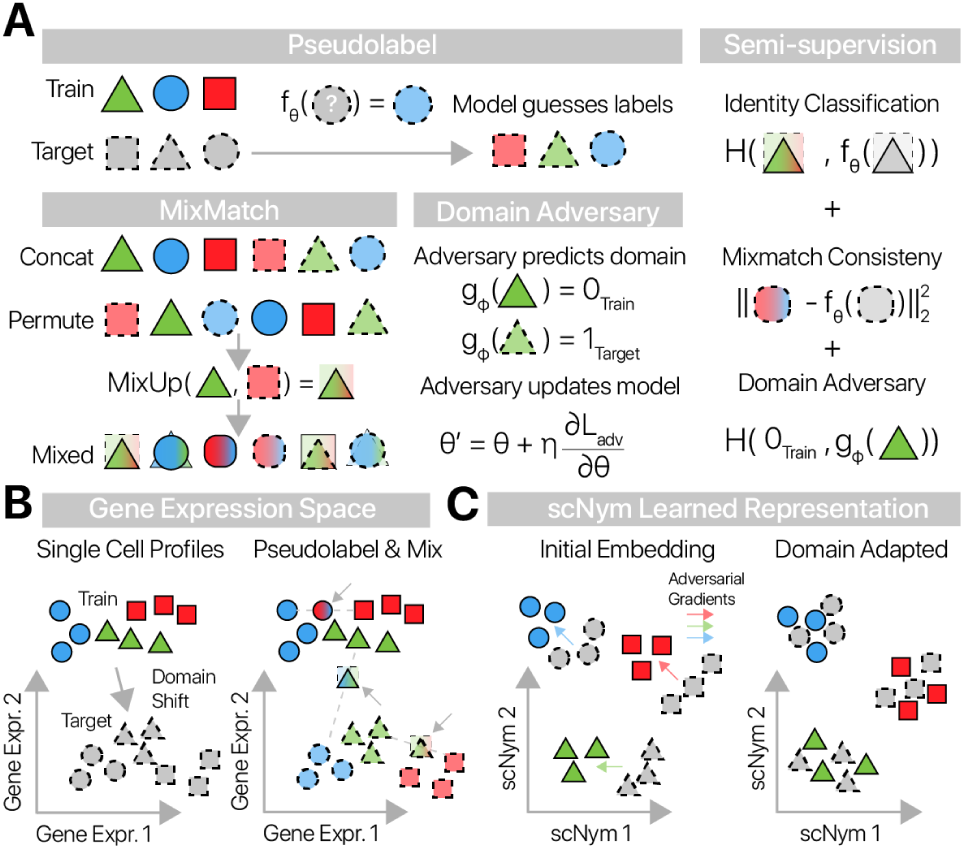
scNym combines semi-supervised and adversarial training to learn performant single cell classifiers. Graphical depiction of the scNym training procedure. **(A)** Pseudolabels are generated for each observation in the unlabeled target data using model predictions and augmented with MixUp. An adversary is also trained to discriminate training and target observations. We train model parameters using a combination of supervised classification, interpolation consistency, and adversarial objectives.**(B)** Training and target cell profiles are separated by a domain shift in gene expression space. scNym pseudolabels target profiles and generates mixed cell profiles (arrows) by randomly pairing cells. **(C)** scNym models learn a rich representation of cell state in a hidden embedding layer. Train and target cell profiles initially segregate in this representation. During training, adversarial gradients (colored arrows) encourage cells of the same type to mix in the scNym embedding.

The scNym classifier learns a rich representation of cell identity in the hidden neural network layers to enable cell type classification. Alongside the cell type classifier, we train an adversarial model to predict the domain-of-origin for each cell (e.g. training set, target set) from this learned embedding. We train the scNym classifier to compete with this adversary, updating the classifier’s embedding to make domain prediction more difficult at each iteration. This adversarial training procedure encourages the classification model to learn a domain adapted embedding of the training and target datasets that improves classification performance (Fig. 1C).

Our model requires no prior manual specification of marker genes, but rather learns relevant gene expression features from the data. After training, scNym predictions provide a probability distribution across all cell types in the training set for target cell profiles. The probability mass on the most likely cell type serves as an intuitive metric of classification confidence, allowing a domain expert to prioritize model decisions for review.

### scNym transfers cell annotations across biological conditions

We evaluated the performance of scNym transferring cell identity annotations in seven distinct tasks. These tasks were chosen to capture diverse kinds of technological and biological variation that complicate annotation transfer. Each task represents a true cell type transfer across different experiments, in contrast to some efforts that report within-experiment hold-out accuracy.

We first evaluated cell type annotation transfer between animals of different ages. We trained scNym models on cells from young rats (5 months old) from the Rat Aging Cell Atlas [4] and predicted on cells from aged rats (27 months old, Fig. 2A, Methods). We found that predictions from our scNym model trained on young cells largely matched the ground truth annotations (92.2% accurate) on aged cells (Fig. 2B, C).

**Figure 2:**
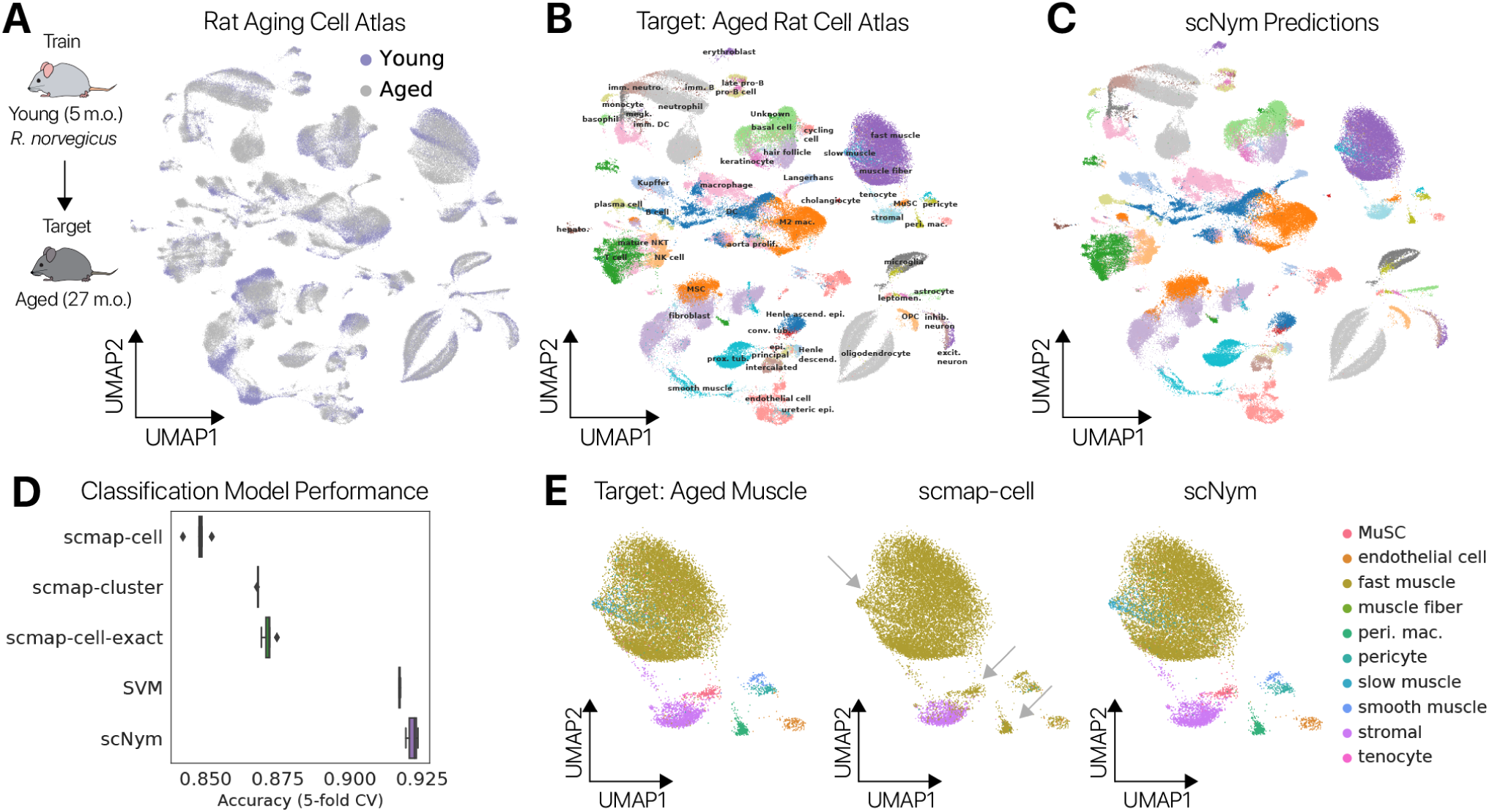
scNym transfers cell identity annotations between young and aged rat cells. **(A)** Young and aged cells from a rat aging cell atlas displayed in a UMAP projection. Some cell types show a domain shift between young and aged cells. scNym models were trained on young cells in the atlas and used to predict labels for aged cells. **(B)** Ground truth cell type annotations for the aged cells of the Rat Aging Cell Atlas shown in a UMAP projection. **(C)** scNym predicted cell types in the target dataset. scNym predictions match ground truth annotation in the majority (>90%) of cases. **(D)** Accuracy scores for scNym models and other state-of-the-art classification models. We find that scNym yields significantly higher accuracy scores than baseline methods (*p* < 0.01, Wilcoxon Rank Sums). Note: multiple existing methods could not complete this task. **(E)** Skeletal muscle cells labeled with ground truth annotations (left), scmap-cell predictions (center), and scNym predictions (right) are displayed in a UMAP projection. scNym accurately predicts multiple cell types that are confused by scmap-cell (arrows).

We compared scNym performance on this task to state-of-the-art single cell identity annotation methods [6, 7, 8, 9, 10] (Methods). scNym produced significantly improved labels over these methods, some of which could not complete this large task on our hardware (256GB RAM) (Wilcoxon Rank Sums, *p* < 0.01, Fig. 2D, Table 1). We found that some of the largest differences in accuracy between scNym and the commonly used scmap-cell method were in the skeletal muscle. scNym models accurately classified multiple cell types in the muscle that were confused by scmap-cell (Fig. 2E), demonstrating that the increased accuracy of scNym is meaningful for downstream analyses.

**Table 1:**
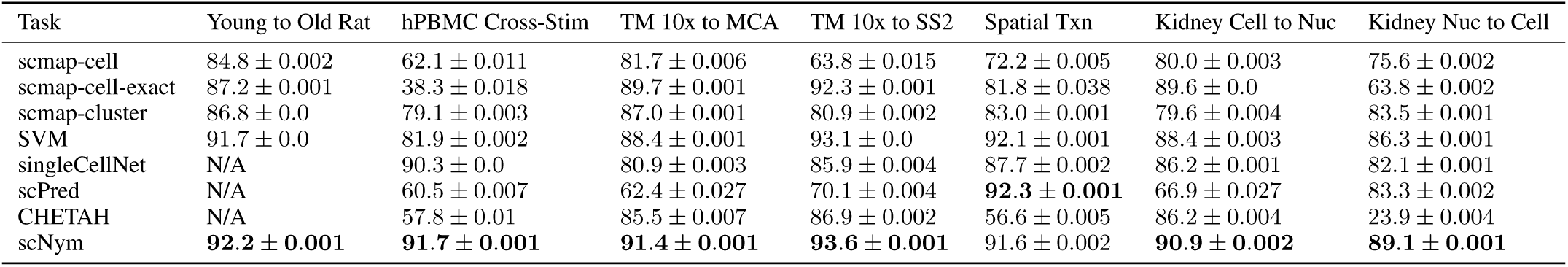
Comparison of model performance across tasks. Mean accuracy *±* standard error across 5-fold training split is reported. Bold text marks best models per task (*p* < 0.05, Rank Sums test). N/A indicates that a model could not complete the task on our hardware (256 GB of RAM). scNym is the only model to achieve high performance (> 85% accuracy) on all tasks.

We next tested the ability of scNym to classify cell identities after perturbation. We trained on unstimulated human peripheral blood mononuclear cells (PBMCs) and predicted on PBMCs after stimulation with IFN*β* [21]. scNym achieved high accuracy (> 91%), superior to baseline methods (Fig. 3A, Table 1). The common scmap-cluster method frequently confused monocyte subtypes, while scNym did not (Fig. 3B).

**Figure 3:**
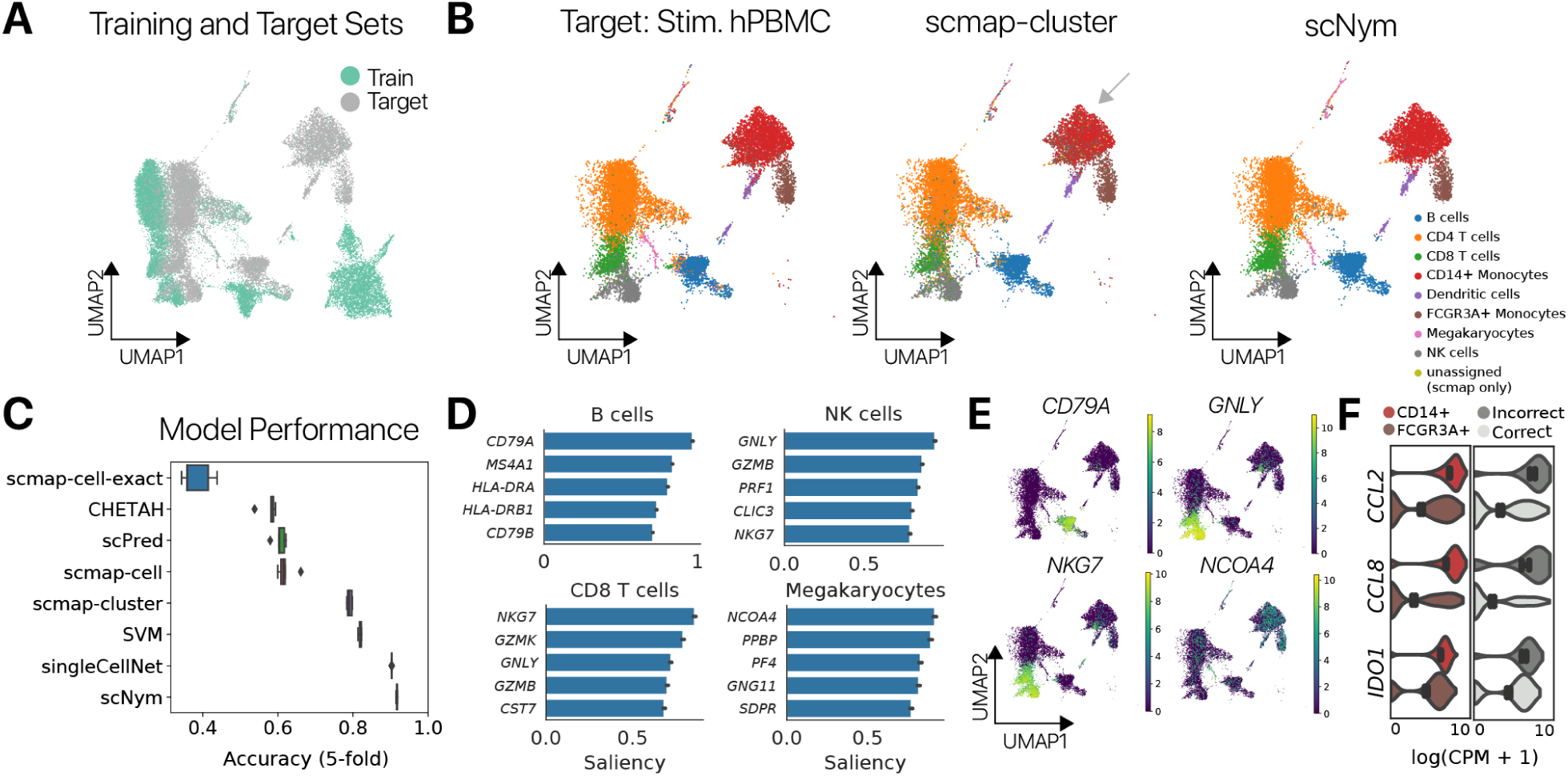
scNym transfers annotations from unstimulated immune cells to stimulated immune cells. **(A)** UMAP projection of unstimulated PBMC training data and stimulated PBMC target data with stimulation condition labels. **(B)** UMAP projections of ground truth cell type labels (left), scmap-cluster predictions (center), and scNym predictions (right). scNym provides consistent annotations for both CD14+ and FCGR3A + monocytes. scmap-cluster appears to confuse these populations (arrow). **(C)** Classification accuracy for scNym and baseline cell identity classification methods. scNym is significantly more accurate than other approaches (*p* < 0.01, Wilcoxon Rank Sums). **(D)** Saliency analysis reveals genes that drive correct classification decisions. We recover known marker genes of many cell types (e.g. *CD79A* for B cells, *PPBP* for magekaryocytes). **(E)** Cell type specificity of the top salient genes in a UMAP projection of gene expression (log normalized counts per million). **(F)** Saliency analysis reveals genes that drive incorrect classification of a some FCGR3A+ monocytes as CD14+ monocytes. Several of the top 15 salient genes for misclassification are CD14+ markers that are upregulated in incorrectly classified FCGR3A+ cells.

### scNym models learn biologically meaningful cell type representations

To interpret the classification decisions of our scNym mod-els, we developed saliency analysis tools to identify genes that influence model decisions (Methods). We found that salient genes included known markers of specific cell types such as *CD79A* for B cells and *GNLY* for NK cells. Saliency analysis also revealed specific cell type marker genes that may not have been selected *a priori*, such as *NCOA4* for megakaryocytes (Fig. 3D, Fig. S1). This result demonstrates that our models are learning biologically meaningful representations.

We also used saliency analysis to understand why the scNym model misclassified some FCGR3A+ monocytes as CD14+ monocytes (Methods). This analysis revealed genes driving these incorrect classifications (Fig. 3F), including some CD14+ monocyte marker genes that are elevated in a subset of FCGR3A+ monocytes (Fig. 3G). Domain experts may use saliency analysis to understand and review model decisions for ambiguous cells.

### scNym transfers annotations across single cell sequencing technologies

To evaluate the ability of scNym models to transfer labels across different experimental technologies, we trained scNym models on single cell profiles from ten mouse tissues in the “Tabula Muris” captured using the 10x Chromium technology and predicted labels for cells from the same experiment captured using Smart-Seq2 [22]. We found that scNym predictions were highly accurate (> 90%) and superior to baseline methods (Fig. S3A, B, C). scNym models accurately classified monocyte subtypes, while baseline methods frequently confused these cells (Fig. S3D, E).

In a second cross-technology task, we trained scNym on mouse lung data from the Tabula Muris and predicted on lung data from the “Mouse Cell Atlas,” a separate experimental effort that used the Microwell-Seq technology [23]. We found that scNym yielded high classification accuracy (> 90%), superior to baseline methods, despite experimental batch effects and differences in the sequencing technologies (Fig. S4). We also trained scNym models to transfer regional identity annotations in spatial transcriptomics data and found performance competitive with baseline methods (Fig. S5). Together, these results demonstrate that scNym models can transfer cell type annotations across technologies and experimental environments with high performance.

### Multi-domain training allows integration of multiple reference datasets

The number of public single cell datasets is increasing rapidly [24]. Integrating information across multiple reference datasets may improve annotation transfer performance on challenging tasks. The domain adversarial training framework in scNym naturally extends to training across multiple reference datasets. We hypothesized that a multi-domain training approach would allow for more general representations that improve annotation transfer. To test this hypothesis, we evaluated the performance of scNym to transfer annotations between single cell and single nucleus RNA-seq experiments in the mouse kidney. These data contained six different single cell preparation methods and three different single nucleus methods, capturing a range of technical variation in nine distinct domains [25](Fig. 4A, B).

scNym models achieved significantly higher accuracy than baseline methods transferring labels from single cell to single nucleus experiments using multi-domain training. This result was also achieved for the inverse transfer task, transferring annotations from single nucleus to single cell experiments (Fig. 4C, Table 1). We found that scNym models offered more accurate annotations for multiple cell types in the cell to nucleus transfer task, including mesangial cells and tubule cell types (Fig. 4D, E). These improved annotations highlight that the performance advantages of scNym are meaningful for downstream analysis and biological interpretation. We found that multi-domain scNym models achieved higher accuracy than any single domain model on both tasks and effectively synthesized information from high- and low-performance single domain training sets (Fig. 4F, Fig. S6).

**Figure 4:**
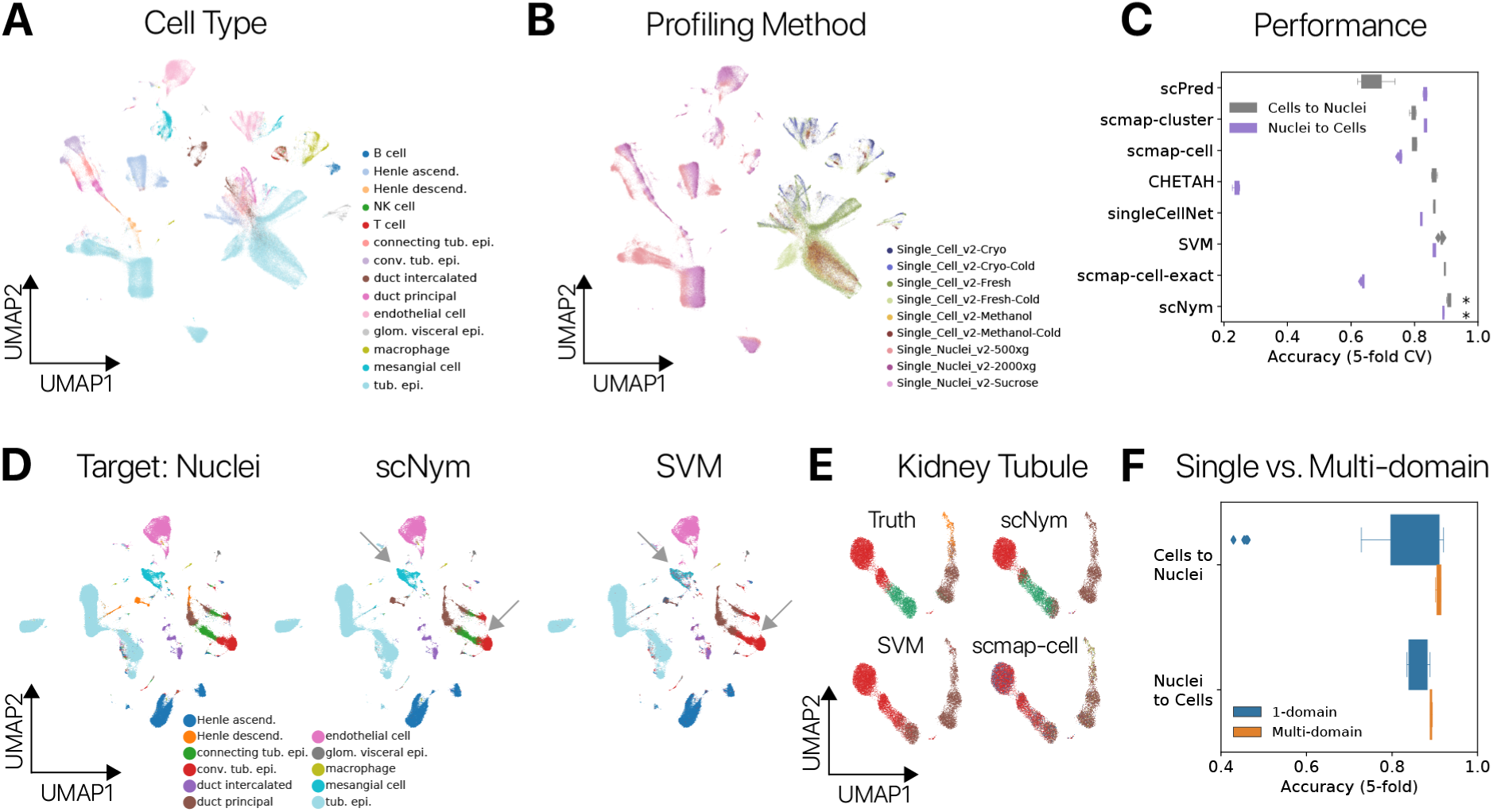
Multi-domain training improves cross-technology annotation transfer in the mouse kidney. **(A)** Cell type and **(B)** sequencing protocol annotations overlaid on a UMAP projection of single cell and single nucleus RNA-seq profiles from the mouse kidney. Each protocol in **(B)** represents a unique training domain that captures technical variation. **(C)** Performance of scNym and baseline approaches on single cell to nucleus and single nucleus to cell annotation transfer. scNym offers significantly superior performance (*: Wilcoxon Rank Sum, *p* < 0.01) on both tasks. **(D)** Ground truth annotations and model predictions for single nucleus target data in the single cell to nucleus transfer task. scNym achieves more accurate labeling of mesangial cells and tubule cell types (arrows). **(E)** Kidney tubule cells from **(D)** visualized independently with true and predicted labels. scNym offers the closest match to true annotations. All methods make notable errors on this difficult task. **(F)** Comparison of scNym performance when trained on individual training datasets (1-domain) vs. multi-domain training across all available datasets. We found that multi-domain training improves performance on both the cells to nuclei and nuclei to cells transfer tasks (Wilcoxon Rank Sums, *p* = 0.073 and *p* < 0.01 respectively).

### scNym confidence scores enable expert review and new cell type discoveries

Calibrated predictions, in which the classification probabil-ity returned by the model precisely reflects the probability it is correct, enable more effective interaction of the hu-man researcher with the model output. We investigated scNym calibration by comparing the prediction confidence scores to prediction accuracy (Methods). We found that semi-supervised adversarial training improved model cal-ibration, such that high confidence predictions are more likely to be correct (Fig. 5A, B; Fig. S7A, B; Fig. S8).

**Figure 5:**
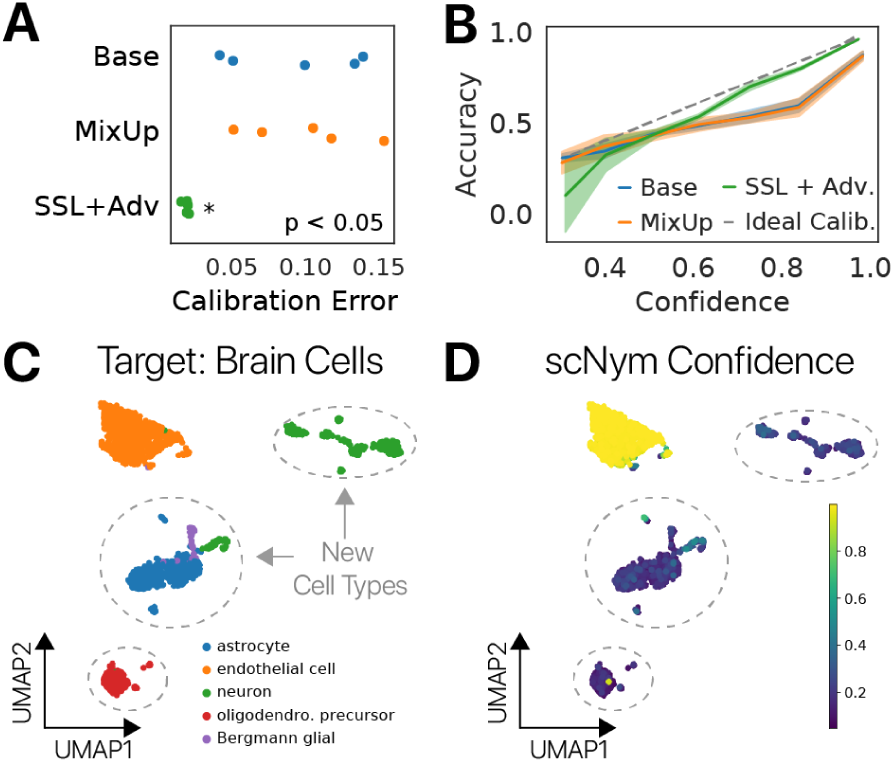
scNym confidence scores highlight new cell type discoveries. **(A)** Calibration error for scNym models trained on the human PBMC cross-stimulation task. We found that semi-supervised and adversarial training signifi-cantly reduced calibration error relative to models trained with only supervised methods (Base, MixUp). **(B)** Calibration curves capturing the relationship between model confidence and empirical accuracy for models in **(A). (C)** scNym models were trained to transfer annotations from a mouse atlas without brain cell types to data from mouse brain tissue. We desire a model that provides low confi-dence scores to the new cell types and high confidence scores for endothelial cells seen in other tissues. **(D)** sc-Nym confidence scores for target brain cells. New cell types receive low confidence scores as desired (dashed outlines).

scNym confidence scores can therefore be used to highlight cells that may benefit from manual review (Fig. S7C, Fig. S8B), further improving the annotation exercise when it contains a domain expert in the loop.

scNym confidence scores can also highlight new, unseen cell types in the target dataset using an optional pseudolabel thresholding procedure during training, inspired by FixMatch [26] (Methods). The semi-supervised and adversarial components of scNym encourage the model to find a matching identity for cells in the target dataset. Pseudolabel thresholding allows scNym to exclude cells with low confidence pseudolabels from the semi-supervised and adversarial components of training, stopping these components from mismatching unseen cell types and resulting in correctly uncertain predictions.

To test this approach, we simulated two experiments where we “discover” multiple cell types by predicting annotations on the Tabula Muris brain cell data using models trained on non-brain tissues (Fig. 5A, B; Methods). We first used pre-trained scNym models to predict labels for new cell types not present in the original training or target sets, and scNym correctly marked these cells with low confidence scores (Fig. S9). In the second experiment, we included new cell types in the target set during training and found that scNym models with pseudolabel thresholding correctly provided low confidence scores to new cell types, highlighting these cells as potential cell type discoveries for manual inspection. (Fig. 5C, D; Fig. S10).

### Semi-supervised adversarial training improves annotation transfer

We ablated different components of our scNym model to determine which features were responsible for high performance. We found that semi-supervision with MixMatch and training with a domain adversary improved model performance across multiple tasks (Fig. 6B, Fig. S2). We hypothesized that scNym models might benefit from domain adaptation through the adversarial model. We found that training and target domains were significantly more mixed in scNym embeddings, supporting this hypothesis (Fig. S11). These results suggest that semi-supervision and adversarial training improve the accuracy of cell type classifications.

**Figure 6:**
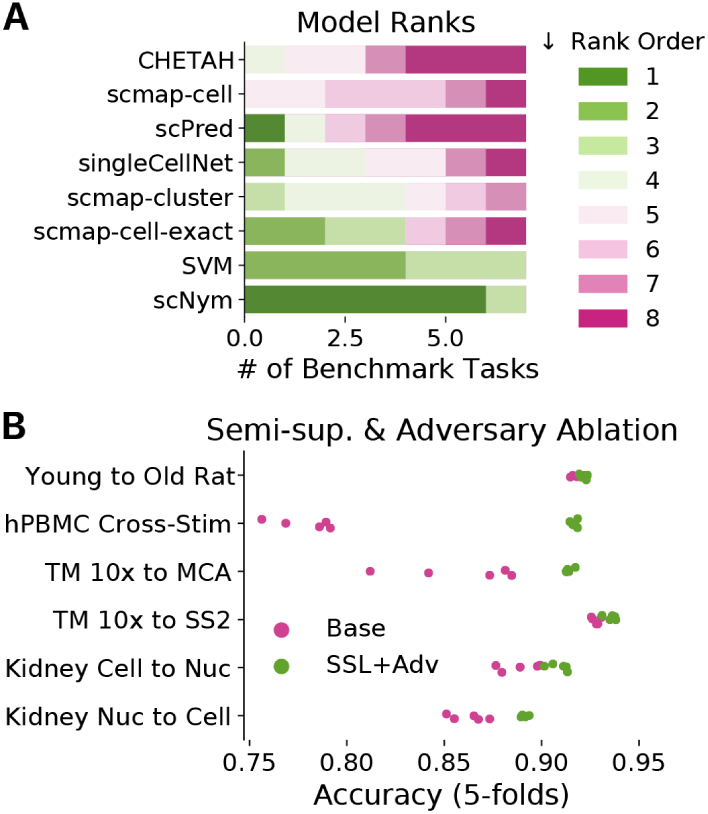
Comparison of semi-supervised scNym to other single cell classification methods and ablated sc-Nym variants. **(A)** We assign each method a rank order (Rank 1 is best) based on performance for each bench-mark task. scNym is superior to all benchmark methods on 6/7 tasks, and highly ranked on the seventh. A support vector machine (SVM) baseline is the next best method, consistent with a previous benchmarking study. **(B)** Ablation experiments comparing simplified supervised scNym models (Base) against the full scNym model with semi-supervised and adversarial training (SSL + Adv.). We found that semi-supervised and adversarial training significantly improved scNym performance across diverse tasks (all tasks, Wilcoxon Rank Sum, *p* < 0.05).

## Discussion

Single cell genomics experiments have become more accessible due to commercial technologies, enabling a rapid increase in the use of these methods [27]. Cell identity annotation is an essential step in the analysis of these exper-iments, motivating the development of high performance, automated annotation methods that can take advantage of diverse datasets. Here, we introduced a semi-supervised adversarial neural network model that learns to transfer annotations from one experiment to another, taking advantage of information in both labeled training sets and an unlabeled target dataset. Our benchmark experiments demonstrate that these scNym models provide high performance across a range of cell identity classification tasks, including cross-age, cross-perturbation, and cross-technology scenarios. scNym performs better than six state-of-the-art baseline methods in these varied conditions and is the only method with high performance (>90% accuracy on all tasks, Fig. 6, Table 1).

The key idea that differentiates scNym from previous cell classification approaches is the use of semi-supervised [16] and adversarial training [17] to extract information from the unlabeled, target experiment we wish to annotate. Through ablation experiments, we showed that these training strategies improve the performance of our models. Performance improvements were most pronounced when there were large, systematic differences between the training and target datasets (Fig. 3). Semi-supervision and adversarial training also allow scNym to integrate information across multiple training and target datasets, improving performance (Fig. 4). As large scale single cell perturbation experiments become more common [28, 29] and multiple cell atlases are released for common model systems, our method’s ability to adapt across distinct biological and technical conditions will only increase in value.

Most downstream biological analyses rely upon cell identity annotations, so it is important that researchers are able to interpret the molecular features that drive model decisions. We showed that backpropogation-based saliency analysis methods are able to recover specific cell type markers, confirming that scNym models learn interpretable, biologically relevant features of cell type. In future work, we hope to extend upon these interpretability methods to infer perturbations that alter cell identity programs using rich representations learned by scNym. We aim to enable widespread use of these methods via the release of their open source implementations, tutorials, and pre-trained models for mouse, rat, and human cell types available from https://www.github.com/calico/scnym.

## Methods

### scNym Model

Our scNym model *f*_*θ*_ consists of a neural network with an input layer, two hidden layers, each with 256 nodes, and an output layer with a node for each class. The first three layers are paired with batch normalization, rectified linear unit activation, [30], and dropout [31]. The final layer is paired with a softmax activation to transform raw outputs of the neural network into a vector of class probabilities. The model maps cell profile vectors *x* to probability distributions *p*(*y*| *x*) over cell identity classes *y*.

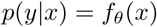

We train scNym models to map cell profiles in a gene expression matrix *x ∈* **X**^Cells*×*Genes^ to paired cell iden-tity annotations *y ∈***y**. Transcript counts in the gene expression matrix are normalized to counts per million (CPM) and log-transformed after addition of a pseudo-count (log(CPM + 1)). During training, we randomly mask 10% of genes in each cell with 0 values.

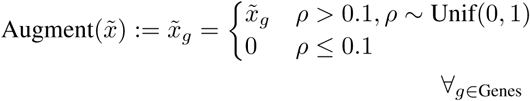

After renormalizing the cell profile, this process generates an augmented cell profile *x* for each raw profile 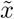

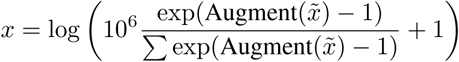

We use the Adadelta adaptive stochastic gradient descent method [32] with an initial learning rate of *η* = 1.0 to update model parameters on minibatches of cells, with batch sizes of 256. We apply a weight decay term of *λ*_WD_ = 10^−4^ for regularization. We train scNym models to minimize a standard cross-entropy loss function for supervised training.

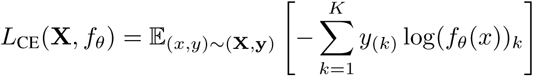

where *y*_(*k*)_ is an indicator variable for the membership of *x* in class *k*, and *k ∈ K* represent class indicators.

We fit all scNym models for a maximum of 400 epochs and selected the optimal set of weights using early stopping on a validation set consisting of 10% of the training data. We initiate early stopping after training has completed at least 5% of the total epochs to avoid premature termination.

Prior to passing each minibatch to the network, we perform dynamic data augmentation with the “MixUp” operation [18]. MixUp computes a weighted average of two samples *x* and *x*^*′*^ where the weights *λ* are randomly sampled from a Beta distribution with a symmetric shape parameter *α*.

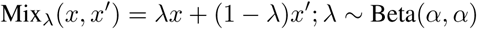

For all experiments here, we set *α* = 0.3 based on performance in the natural image domain [18]. Forcing models to interpolate predictions smoothly between samples shifts the decision boundary away from high-density regions of the input distribution, improving generalization. This procedure has been shown to improve classifier performance on multiple tasks [18]. Model calibration – the correctness of a model’s confidence scores for each class – is generally also improved by this augmentation scheme [19].

### Semi-supervision with MixMatch

We train semi-supervised scNym models using the MixMatch framework [16], treating the target dataset as unlabeled data 𝒰. At each iteration, MixMatch samples mini-batches from both the labeled dataset (**X, y**) ∼ 𝒟 and unlabeled dataset **U** *∼* 𝒰 We generate “pseudolabels” [20] using model predictions for each observation in the unlabeled minibatch. While generating pseudolabels, we run the model in typical inference mode–deactivating dropout layers, not updating batch normalization running statistics, and not propagating gradients. We use the running estimates of feature mean and variance computed during training for batch normalization during pseudolabeling.

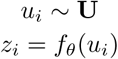

Our procedure differs slightly from standard MixMatch in that we do not average across several augmented versions of unlabeled observations, but rather pseudolabel a single, unmodified observation. This choice was motivated by empirical performance. Augmentation ensembling may be unable to deliver improved accuracy because only simple augmentations are available for gene expression data, relative to the rich set of transformations available for the imaging data originally studied with MixMatch.

We next “sharpen” the pseudolabels using a scaling procedure, often referred to as “temperature scaling” [33, 34].

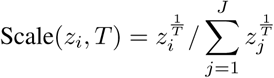

where *z*_*i*_ is a probability vector and *T* is a temperature parameter. Values of *T* > 1 increase the entropy of the probability vector (“smoothing”), while values of *T* < 1 decrease the entropy of the probability vector (“sharpen-ing”). For our experiments, we set *T* = 0.5 such that the transformation acts as a form of entropy minimization on the pseudolabels. This entropy minimization encourages unlabeled examples to belong to one of the described classes.

We then randomly mix each observation in both the labeled and unlabeled minibatches with another observation using MixUp [18]. We allow labeled and unlabeled observations to mix together during this procedure. As in standard MixUp training, we mix both the observed features and the labels/pseudolabels between the two randomly paired observations.

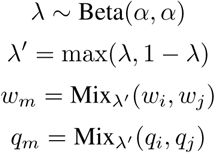

where (*w*_*i*_, *q*_*i*_) is either a labeled observation and ground truth label (*x*_*i*_, *y*_*i*_) or an unlabeled observation and the pseudolabel (*u*_*i*_, *z*_*i*_).

We set the MixUp parameter *λ*^′^ = max(*λ*, 1 − *λ*) such that the first example in the MixUp operation always dominates the resulting mixed sample. This allows us to maintain the labeled or unlabeled identity of each observation in the mixed minibatch. This procedure yields a minibatch **X**^*′*^ of mixed labeled observations and a minibatch **U**^′^ of mixed unlabeled observations.

We introduce a semi-supervised interpolation consistency penalty during training in addition to the standard supervised loss. For observations and pseudolabels in the mixed unlabeled minibatch *U* ^′^, we penalize the mean squared error (MSE) between the mixed pseudo-labels and the model prediction for the mixed observation.

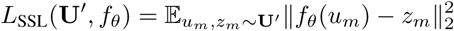

This encourages the model to provide smooth interpolations between observations and their ground truth or pseudo-labels, generalizing the decision boundary of the model. We use MSE on the model predictions and mixed pseudo-label rather than the cross-entropy because it is bounded and therefore less sensitive to incorrect predictions [16].

We weight this unsupervised loss relative to the supervised cross-entropy loss using a weighting function *λ*_SSL_(*t*) → [0, 1].

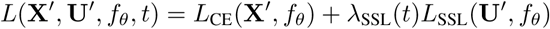

We initialize this coefficient to *λ*_SSL_ = 0 and increase the weight to a final value of *λ*_SSL_ = 1 over 100 epochs using a sigmoid schedule. We set *λ*_SSL_ based on recommendations from the MixMatch authors. We adopt a sigmoid scaling function that is common in the semi-supervised learning field [15].

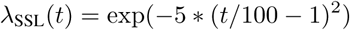

### Domain Adaptation with Domain Adversarial Networks

We use domain adversarial networks (DAN) as an additional approach to incorporate information from the target dataset during training [17]. DANs are one of a family of domain adaptation approaches that attempt to transfer labels from a training set to a target set, where the target distribution is systematically different than the training distribution. We can readily appreciate the application to single cell identity classification, as the training and target domains are likely to differ across technologies, conditions, batches, and laboratories.

The DAN method encourages the classification model to embed cells from the training and target dataset with similar coordinates, such that training and target datasets are well-mixed in the embedding. By encouraging the training and target dataset to be well-mixed, we take advantage of the inductive bias that cell identity classes in each dataset are similar, despite technical variation or differences in conditions. This bias allows the model to adapt across domains, producing an embedding where training and test data are well-mixed.

If cell type proportions differ dramatically between training and target domains, this inductive bias may be ill-suited and the domain adversary could conceivably harm performance. For this reason, we use a low weight for the adversarial penalty. Empirically, we find that the adversary improves performance even when the cell type distributions are imbalanced. For example, we find that in our Tabula Muris 10x Chromium to Mouse Cell Atlas transfer experiment, the adversary improves performance even though the cell type distributions differ drastically (Fig. S4).

We introduce this technique into scNym by adding an adversarial domain classification network *g*_*ϕ*_. We implement *g*_*ϕ*_ as a two-layer neural network with a single hidden layer of 256 units and a rectified linear unit activation, followed by a classification layer with two outputs and a softmax activation. This adversary attempts to predict the domain of origin *d* from the penultimate classifier embedding *v* of each observation. For each forward pass, it outputs a probability vector 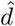 estimating the likelihood the observation came from the training or test domain.

We assign a domain label *d* to each molecular profile based on the experiment of origin. For experiments with only a single training dataset and single target dataset, we set cells from the training dataset to *d* = 0 and set *d* = 1 for cells from the target dataset. For multi-domain training experiments in the mouse kidney, we assigned a unique domain label to each sequencing protocol in the training and target datasets (Fig. 4B). We one-hot encode all domain labels.

During training, we pass a minibatch of labeled observa-tions *x ∈***X** and unlabeled observations *u ∈***U** through the domain adversary to predict domain labels.

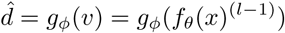

where 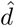 is the domain probability vector and *v* = *f*_*θ*_(*x*)^(*l*−1)^ denotes the embedding of *x* from the penultimate layer of the classification model *f*_*θ*_.

We fit the adversary using a multi-class cross-entropy loss:

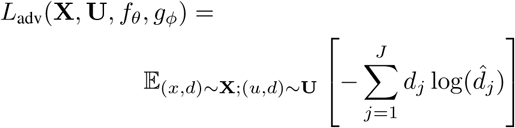

where *j ∈ J* are domain indicator indices and *J* is the set of source domains.

To make use of the adversary for training the classification model, we use the “gradient reversal” trick at each back-ward pass. We update the parameters *ϕ* of the adversary using the gradients computed during a backward pass from *L*_adv_. At each backward pass, this optimization improves the adversarial domain classifier. Our update rule for the parameters *ϕ* is therefore a standard gradient descent:

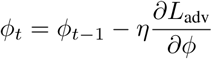

where *η* is the learning rate.

We update the parameters *θ* of the classification model using the *inverse* of the gradients computed during a backward pass from *L*_adv_. Using the inverse gradients encourages the classification model *f*_*θ*_ to generate an embedding where it is difficult for the adversary to predict the domain. Our update rule for the classification model parameters therefore becomes:

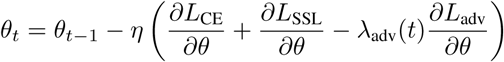

We increase the weight of the adversary gradients from *λ*_adv_ → [0, 0.1] over the course of 20 epochs during training using the sigmoid schedule above. By scaling the adversary’s gradients, we allow both the classification model *f*_*θ*_ and the adversary *g*_*ϕ*_ to find an approximate fit before introducing the adversarial penalty. We scale the adversarial *gradients* flowing to *θ*, rather than the adversarial loss term, so that full magnitude gradients are used to train a robust adversary *g*_*ϕ*_.

We set *λ*_adv_ based on the authors’ experiments in the natural image domain [17]. This setting appears sufficient to promote domain adaptation (Fig. S11) while also training a robust classifier (Table 1). The much larger weight of the supervised classification loss relative to the adversarial loss leads to trained models where domain classification accuracy is very high in certain regions, in which reducing it would require collapsing cell types at the expense of the supervised loss (Fig. S12). We found that increasing the weight eventually led the model to pathologically collapse cell types in the embedding (Fig. S13).

Incorporating both MixMatch and the domain adversary, our full loss term becomes:

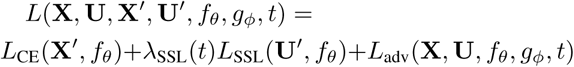

### Pseudolabel Thresholding for New Cell Type Discovery

Entropy minimization and domain adversarial training enforce an inductive bias that all cells in the target dataset belong to a class in the training dataset. For many cell type classification tasks, this assumption is valid and useful. However, it is violated in the case where new, unseen cell types are present in the target dataset. We introduce an alternative training configuration to allow for quantitative identification of new cell types in these instances.

We first adopt a notion of “pseudolabel confidence thresholding” introduced in the FixMatch method [26]. We set a minimum pseudolabel confidence *τ* = 0.9 to identify unlabeled observations with high and low confidence pseudolabels, marked by a binary indicator *c*_*i*_ → *{*0, 1*}*, where *c*_*i*_ = 1 is a high confidence pseudolabel. We determine pseudolabel confidence *before* entropy minimization (label sharpening) is applied. We chose *τ* based on performance in other problem domains [26].

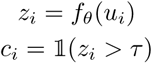

where 𝕝 (*·*) is the indicator function.

We have observed that new cell types will receive low confidence pseudolabels, as they do not closely resemble any of the classes in the training set (Fig. S9). We wish to exclude these low confidence pseudolabels from our entropy minimization and domain adversarial training procedures, as these methods both encourage the classifier to provide more confident labels from the training dataset for unlabeled examples. In the case of new cell types, these procedures may pathologically cause the model to label new cell types with incorrect but high confidence scores for the nearest cell type in the training set.

Pseudolabels influence the model parameters *θ* through each of the three components of our loss – supervised, semi-supervised interpolation consistency, and adversarial. We make two modifications to the training procedure to prevent low confidence pseudolabels from contributing to any component of the loss.

First, we use only high confidence pseudolabels in the MixUp operation of the MixMatch procedure. This prevents low confidence pseudolabels from contributing to the supervised classification loss through MixUp with labeled samples. It also prevents low confidence pseudolabels from mixing with high confidence pseudolabels, allowing us to compute the semi-supervised loss exclusively on high confidence pseudolabels.

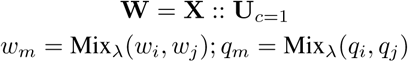

where **W** is a row-wise concatenation of the labeled mini-batch and confident pseudolabeled minibatch and **U**_*c*=1_ are the observations in **U** with confident pseudolabels.

We scale the interpolation consistency loss based on the fraction of confident pseudolabels as in FixMatch [26]. The semi-supervision penalty therefore scales with the number of high confidence observations available.

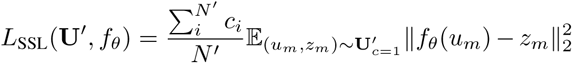

where *N* ^′^is the number of elements in the unlabeled mini-batch.

Second, we use only unlabeled examples with high confidence pseudolabels to train the domain adversary. This prevents unlabeled examples with low confidence pseudolabels from contributing to the adversarial loss. These low confidence unlabeled examples can therefore occupy a unique region in the model embedding, even if they are easily discriminated from training examples.

Our adversarial loss is then only slightly modified to penalize domain predictions only on confident samples in the pseudolabeled minibatch, **U**_*c*=1_.

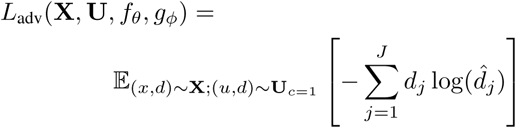

where *j ∈ J* are domain indicator indices and *J* is the set of source domains.

We found that this pseudolabel thresholding configuration option was essential to provide accurate, quantitative information about the presence of new cell types in the target dataset (Fig. S10). However, this option does modestly decrease performance when new cell types are not present. We therefore enable this option when the possibility of new cell types violates the assumption that the training and target data share the same set of cell types. We have provided a simple toggle in our software implementation to allow users to enable or disable this feature.

### scNym Model Embeddings

We generate gene expression embeddings from our scNym model by extracting the activations of the penultimate neural network layer for each cell. We visualize these embeddings using UMAP [35, 36] by constructing a nearest neighbor graph (*k* = 30) in principal component space derived from the penultimate activations. We set `min_dist` = 0.3 for the UMAP minimum distance parameter.

### Entropy of Mixing

We compute the “entropy of mixing” to determine the degree of domain adaptation between training and target datasets in an embedding *X*. The entropy of mixing is defined as the entropy of a vector of class membership in a local neighborhood of the embedding:

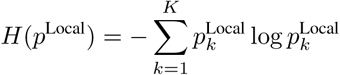

where *p*^Local^ is a vector of class proportions in a local neighborhood and *k ∈ K* are class indices. We compute the entropy of mixing for an embedding *X* by randomly sampling *n* = 1000 cells the embedding, and computing the entropy of mixing on a vector of class proportions for the *k* = 100 nearest neighbors to each point.

### Saliency Analysis

We interpreted the predictions of our scNym models by performing saliency analysis. We compute a saliency score *s* using guided backpropagation to compute gradients on a class probability *f*_*θ*_(*x*)_*k*_ with respect to an input gene expression vector *x*, where *f*_*θ*_ is a trained scNym model [37].

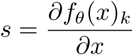

We average these saliency scores across *n*_*s*_ cell input vectors *x* to get class level saliency maps *s*_*k*_, where *n*_*s*_ = min(300, *n*_*k*_) and *n*_*k*_ is the number of cells in the target class.

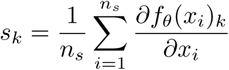

We normalized all values [0, 1] in average saliency maps *s*_*k*_ for presentation. To identify salient genes that drive incorrect classifications, we computed gradients with respect to some class *k* for cells with true class *k*^′^ that were incorrectly classified as class *k*.

### Model Calibration Analysis

We evaluated the calibration of an scNym model by binning all cells in a query set based on the softmax probability of their assigned class – max_*k*_(softmax(*f*_*θ*_(*x*)_*k*_)) – which we term the “confidence score”. We grouped cells into *M* = 10 bins *B*_*m*_ of equal width from [0, 1] and computed the mean accuracy of predictions within each bin.

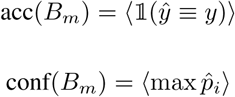

We computed the “expected calibration error” as previously proposed [19].

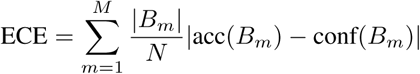

We also computed the “overconfidence error”, which specifically focuses on high confidence but incorrect pre-dictions.

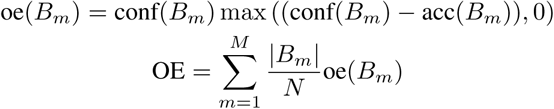

where ⟨*·*⟩ denotes the arithmetic average, *N* is the total number of samples and |*B*_*m*_| is the number of samples in bin *B*_*m*_.

We performed this analysis for each model trained in a 5-fold cross-validation split to estimate variation in calibration for a given model configuration. We evaluated calibrations for baseline scNym models, models with MixUp but not MixMatch, and models with the full MixMatch procedure.

### Dimensionality Reduction

We present single cell experiments using a 2-dimensional representation fit using the UMAP alogirthm [38]. For each experiment, we compute a PCA projection on a set of highly variable genes after log(CPM + 1) normalization. We construct a nearest neighbor graph using first 50 principal components and fit a UMAP projection from this nearest neighbor graph.

### Baseline Methods

As baseline methods, we used seven cell identity classification methods: scmap-cell, scmap-cluster, scmap-cell-exact (scmapcell with exact k-NN search), a linear SVM, scPred, singleCellNet, and CHETAH. For model training, we split data into 5-folds and trained five separate models, each using 4 folds for training and validation data. This allows us to assess variation in model performance as a function of changes in the training data. All models, including scNym, were trained on the same 5-fold splits to ensure equitable access to information.

We applied all baseline methods to all benchmarking tasks. If a method could not complete the task given 256 GB of RAM and 8 CPU cores, we report the accuracy for that method as “Undetermined.” Only scNym models required GPU resources. We trained models on Nvidia K80, GTX1080ti, Titan RTX, or RTX 8000 GPUs, using only a single GPU per model.

### scmap-cluster and scmap-cell

scmap-cluster models were trained using the 1,000 most variable genes selected using “M3Drop” [6, 39]. scmap-cell models were trained with *k* = 10 nearest neighbors and the 1,000 most variable genes selected using “M3Drop” using cosine similarities as a distance metric. The scmap-cell nearest neighbor index was constructed using the default number of subcentroids, 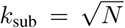 where N is the number of cells. We set scmap hyperparameters to maximize performance based on optimizations in the original publication [6]. We set the minimum similarity parameter threshold = 0:0 to allow scmap methods to predict a class for all cells. However, we found that even with this setting, a small number of cells were given the “unassigned” label. To allow for a more direct comparison, we manually extracted the nearest neighbor graph and called cell types for cells predicted as “unassigned” by scmap-cell.

### scmap-cell-exact

We also implemented a classifier based on scmap-cell with an exact *k*-nearest neighbors search algorithm, rather than the approximate search in the original scmap-cell implementation. We refer to this baseline as “scmap-cell-exact”. We implemented the scmap-cell-exact classifier using scikit-learn “KNeighborsClassifier” with a cosine distance metric. We fit scmap-cell-exact models on the top 1,000 most variable genes selected using the M3Drop procedure with *k* = 10 nearest neighbors, mirroring the settings for the original scmap-cell implementation.

### SVM

Our SVM baseline models were trained using the scikit-learn “LinearSVC” implementation with an *l*_2_ reg-ularization term weighted with *λ*_reg_ = 1.. Models were fit using a squared hinge loss objective and used the “one-versus-rest” multiclass training strategy. This approach and hyperparameter set were suggested as the result of a single-domain benchmarking study [9].

### scPred

scPred models [7] were trained per the authors recommendations, with one exception. We set threshold = 0 so that scPred predicted a class for all observations. This allowed us to compute accuracy scores comparable to other methods, whereas scPred would otherwise report “Undetermined” for many observations.

### singleCellNet

singleCellNet models [8] were trained per the authors recommendations. We used the parameter settings nRand = 70, nTopGenes = 10, nTrees = 1000, nTopGenePairs = 25 as recommended.

### CHETAH

CHETAH models [10] were trained per the authors recommendations. As with scPred models, we set threshold = 0 so that CHETAH models reported a class label for each observation, rather than “Undetermined.” This allowed us to compare CHETAH accuracy to other methods on a fair basis.

### Benchmarking

For the Rat Aging Cell Atlas [4] benchmark, we trained scNym models on single cell RNA-seq data from young, *ad libitum* fed rats (5 months old) and predicted on cells from aged rats (*ad libitum* fed or calorically-restricted). We converted the cell type annotations provided by the authors into “cell ontology” terms for consistency with common nomenclature [40]. We removed genes expressed in < 20 cells to filter uninformative features. We also filtered cells with < 100 or > 5000 expressed genes to remove low-quality profiles and likely multiplets.

For the human PBMC stimulation benchmark, we trained models on unstimulated PBMCs collected from multiple human donors and predicted on IFN*β* stimulated PBMCs collected in the same experiment [21]. Prior to training and prediction, we removed cells without valid cell type labels (“nan”).

For the Tabula Muris cross-technology benchmark, we trained models on Tabula Muris 10x Genomics Chromium platform and predicted on data generated using Smart-Seq2. We limited our analyses to tissues with similar cell type labels between the two technologies: Bladder, Heart and Aorta, Kidney, Limb Muscle, Liver, Lung, Mammary Gland, Spleen, Thymus, and Tongue. We amended the cell ontology class labels of the Smart-seq2 data to match the degenerate names provided in the 10x data. We removed cell types in the Smart-seq2 data that were not present in the 10x Chromium data. We also removed cells with the ambiguous “kidney cell” label from the 10x data which are not found in the Smart-seq2 data. We also corrected labels for a set of “kidney loop of Henle epithelial cells” (*Slc12a1*+, *Umod*+) that were incorrectly labeled as “kidney collecting duct epithelial cells”. Cells were partitioned into clusters using Louvain community detection [41] on a nearest neighbor graph constructed using 50 principal components computed on a set of highly variable genes [42].

For the Mouse Cell Atlas (MCA) [23] benchmark, we trained models on single cell RNA-seq data from lung tissue in the Tabula Muris 10x Chromium data [22] and predicted on MCA lung data. Due to the different cell annotation ontologies in these data sets, we manually reannotated the “Mouse Cell Atlas,” data using the cell ontology class specification so that classes matched the Tabula Muris. During evaluation, we merged “classical monocyte” and “non-classical monocyte” classes predicted by each model to form a single “monocyte” class to match annotations available for the MCA.

For the spatial transcriptomics benchmark, we trained models on spatial transcriptomics from a mouse sagittal-posterior brain section and predicted labels for another brain section. Sagittal-posterior brain section datasets from the 10x Genomics Visium Spatial Transcriptomics demonstration data were downloaded from https://www.10xgenomics.com/resources/datasets/. We manually annotated “region” identities based on the predominant cell type markers in each region. We considered *Neurod1, Cpne7, Ntng1* and *Ddn* to be neuronal markers, *Slc6a11* to be an astrocyte marker, *Plp1* to be an oligodendrocyte marker, and *Car8* and *Sbk1* to be oligodendrocyte precursor cell markers.

For the single cell to single nucleus benchmark in the mouse kidney, we trained scNym models on all single cell data from six unique sequencing protocols and predicted labels for single nuclei from three unique protocols [25]. For the single nucleus to single cell benchmark, we inverted the training and target datasets above to train on the nuclei datasets and predict on the single cell datasets. We set unique domain labels for each protocol during training in both benchmark experiments. To evaluate the impact of multi-domain training, we also trained models on only one single cell or single nucleus protocol using the domains from the opposite technology as target data.

### New Cell Type Discovery Experiments

#### New Cell Type Discovery with Pre-trained Models

We evaluated the ability of scNym to highlight new cell types, unseen in the training data predicting cell type annotations in the Tabula Muris brain cell data (Smart-Seq2) using models trained on 10x data from the ten tissues noted above with the Smart-Seq2 data as corresponding target dataset. No neurons or glia were present in the training or target set for this experiment. This experiment simulates the scenario where a pre-trained model has been fit to transfer across technologies (10x to Smart-Seq2) and is later used to predict cell types in a new tissue, unseen in the original training or target data.

We computed scNym confidence scores for each cell as *c*_*i*_ = max *p*_*i*_, where *p*_*i*_ is the model prediction probability vector for cell *i* as noted above. To highlight potential cell type discoveries, we set a simple threshold on these confidence scores *d*_*i*_ = *c*_*i*_ ≤ 0.5, where *d*_*i*_ *∈*{0, 1} is a binary indicator variable. We found that scNym captured the majority of cells from newly “discovered” types unseen in the training set using this method.

#### New Cell Type Discovery with Semi-supervised Training

We also evaluate the ability of scNym to discover new cell types in a scenario where new cell types are present in the target data used for semisupervised training. We used the same training data and target data as the experiment above, but we now introduce the Tabula Muris brain cell data (Smart-Seq2) into the target dataset during semi-supervised training. We performed this experiment using our default scNym training procedure, as well as the modified new cell type discovery procedure described above.

As above, we conmputed confidence scores for each cell and set a threshold of *d*_*i*_ = *c*_*i*_ *≤* 0.5 to identify potential new cell type discoveries. We found that scNym models trained with the new cell type discovery procedure provided low confidence scores to the new cell types, suitable for identification of these new cells.

## Declarations

### Data and Software Availability

Open source code for our software, pre-processed reference datasets analyzed in this study, and pre-trained models are available in the scNym repository, https://github.com/calico/scnym.

## Author’s Contributions

JCK conceived the study, implemented software, conducted experiments, and wrote the paper. DRK conceived the study and wrote the paper.

## Acknowledgements

We thank Zhenghao Chen, Amoolya H. Singh, and Han Yuan for helpful discussions and comments.

## Supplemental Material

**Figure S1:**
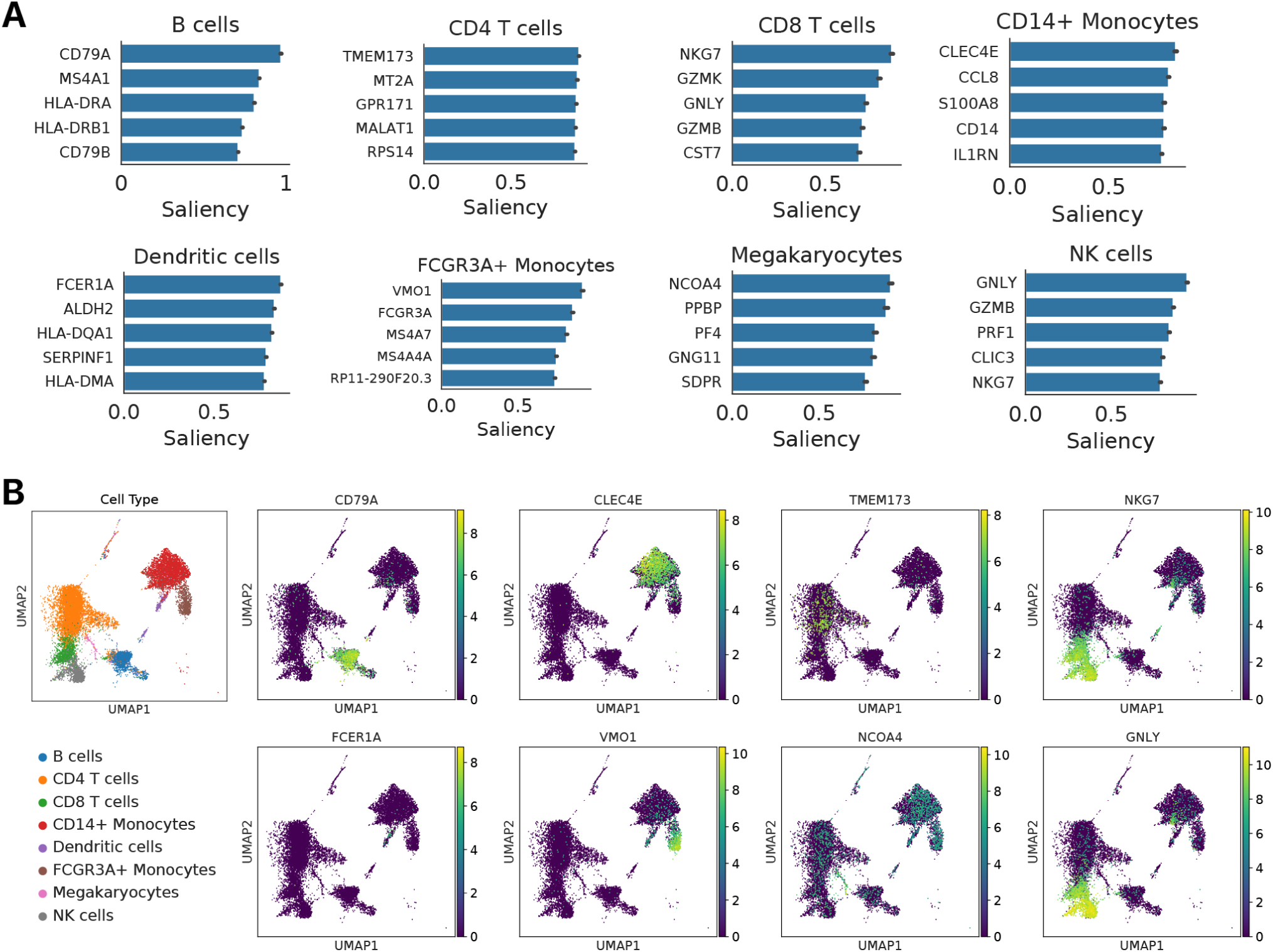
Saliency analysis reveals genes that drive cell type classification decisions. **(A)** Top 5 salient genes for each cell type, computed as the mean across a batch of correctly classified cells. **(B)** Expression of the most salient gene for each cell type in the UMAP embedding.

**Figure S2:**
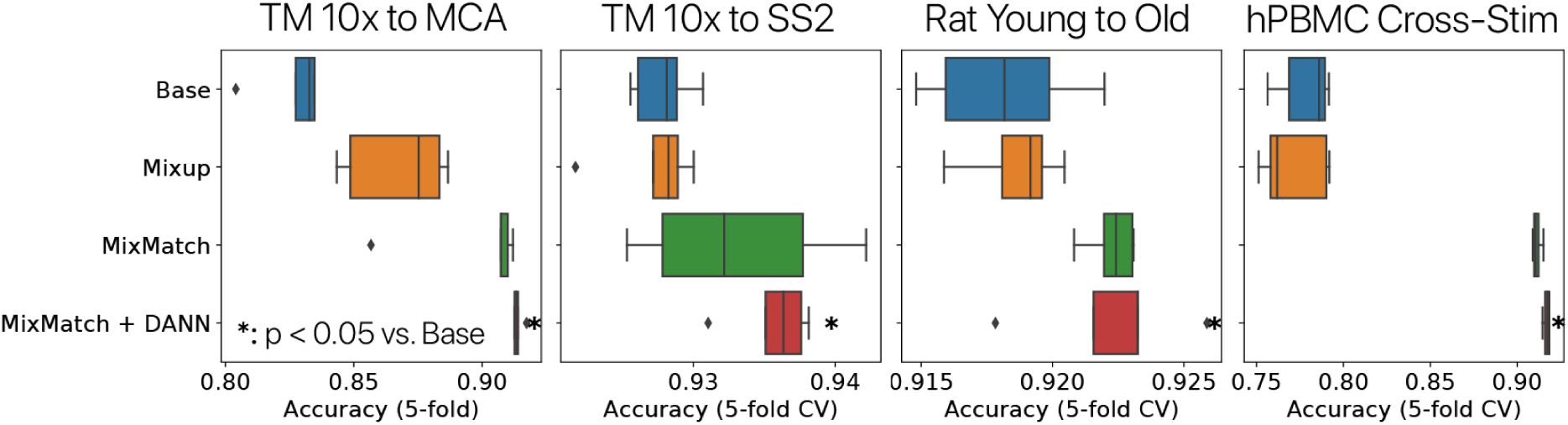
Ablation experiments demonstrate that semi-supervision methods improve scNym performance. We trained scNym models with and without semi-supervision features to complete cell annotation transfer tasks. In each case, we found that inclusion of MixUp augmentations, MixMatch training, and a domain adversary significantly improved scNym performance relative to a base model (*p* < 0.05, Rank Sums test).

**Figure S3:**
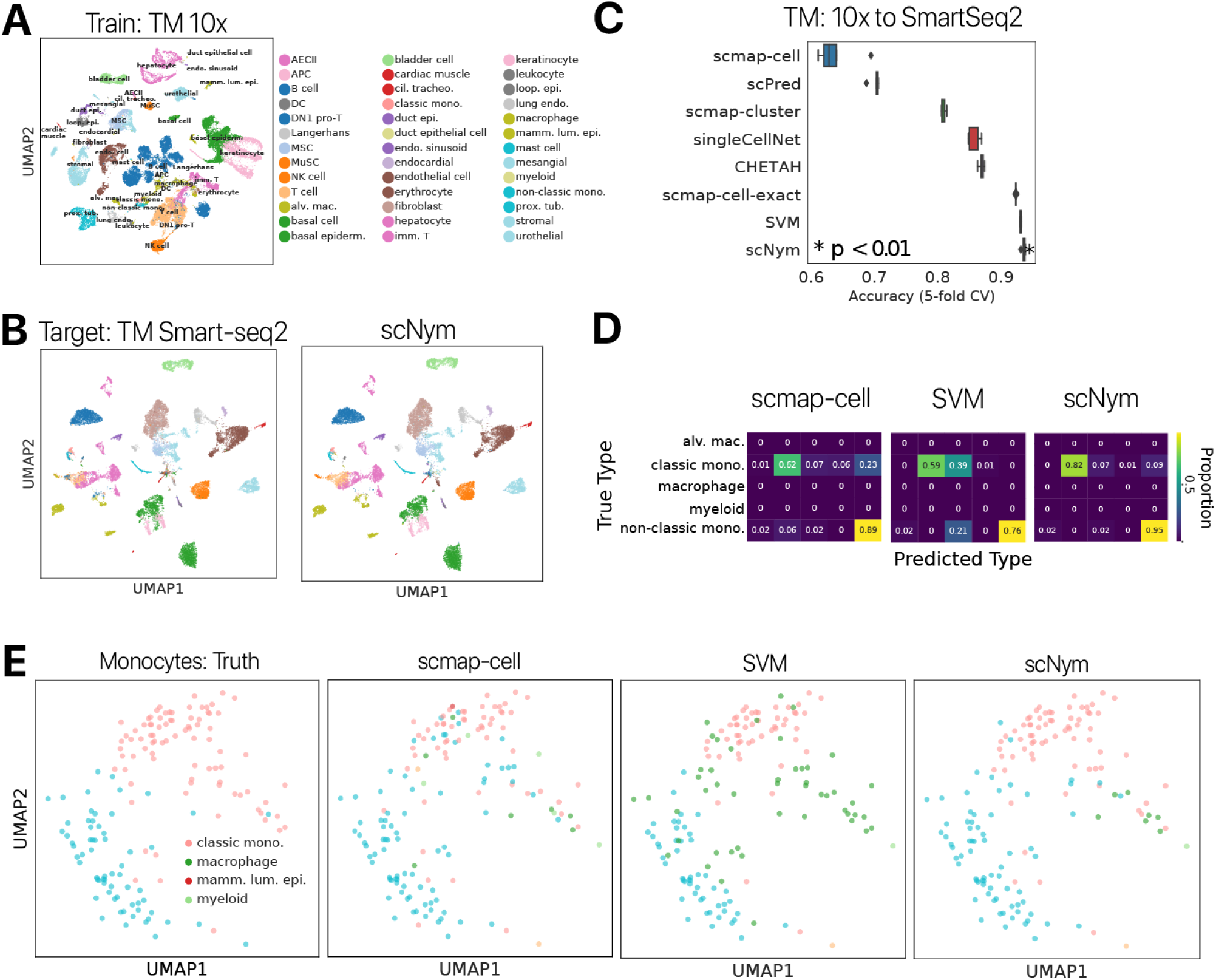
scNym transfers cell identity annotations between 10x and Smart-Seq2 sequencing technologies in the Tabula Muris. **(A)** UMAP projections of ground truth cell type labels in the *Tabula Muris* 10x training data. We limited our analyses to tissues with similar cell type labels between the two technologies: Bladder, Heart and Aorta, Kidney, Limb Muscle, Liver, Lung, Mammary Gland, Spleen, Thymus, and Tongue. **(B)** Ground truth labels and scNym predictions overlaid on a UMAP projection. scNym labels match ground truth for the large majority of cells (>92%). **(C)** Comparison of classification models on the 10x to Smart-Seq2 transfer task. scNym models are significantly more accurate than baseline methods (Wilcoxon Rank Sums on accuracy scores, *p* < 0.01). **(D)** Confusion matrices for scmap-cell, SVM, and scNym models on cells with a ground truth annotation containing “monocyte”. scmap-cell confused classical monocytes and non-classical monocytes, while an SVM confuses monocytes with macrophages. scNym shows less confusion overall. **(E)** Ground truth annotations (left), scmap-cell predictions (left center), SVM predictions (right center) and scNym predictions (right) for monocytes in the Tabula Muris. scNym model predictions more accurately label both types of monocytes.

**Figure S4:**
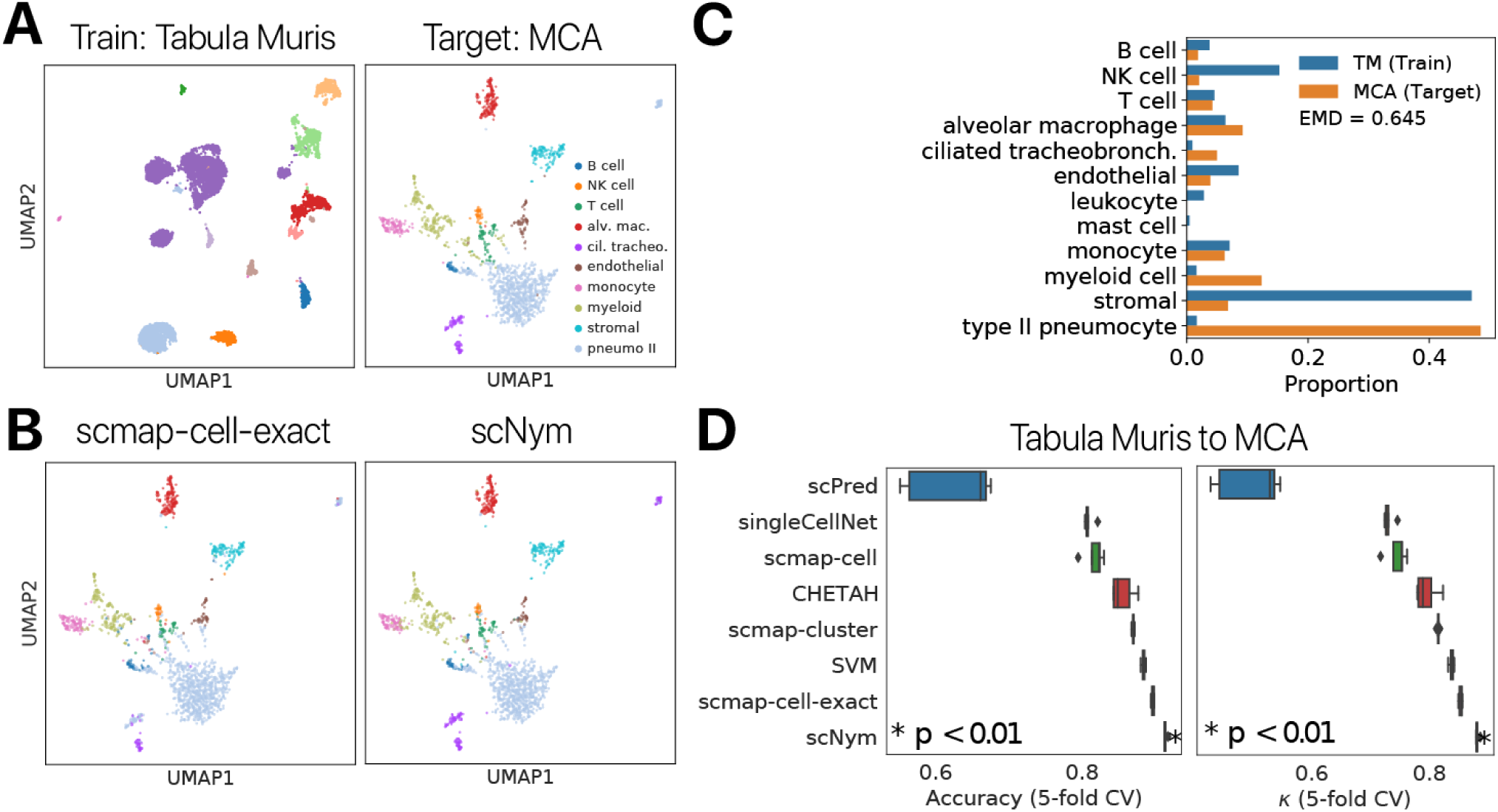
scNym transfers cell identity annotations in the mouse lung between 10x Chromium data from the Tabula Muris and Microwell-seq data from the Mouse Cell Atlas. **(A)** Ground truth annotations for the Tabula Muris training data and MCA target data are displayed in a UMAP projection. **(B)** Predictions from scNym and the next best model scmap-cell-exact are likewise shown in the UMAP projection. scNym predictions are highly concordant with ground truth labels. **(C)** Comparison of cell type distributions between the training set (Tabula Muris, TM) and the target set (Mouse Cell Atlas, MCA). There is a large class imbalance between the two data sets. We quantified this imbalance using the Earth Mover Distance (EMD) and found the EMD to be roughly 0.65, indicating the majority of probability mass would need to shift to match the distributions. Despite this class imbalance, scNym classifiers achieved high performance. **(D)** Accuracy and *κ* score comparison of scNym to seven other cell type classifiers. scNym models show significantly higher performance than the baseline methods (Wilcoxon Rank Sums on accuracy scores, *p* < 0.01).

**Figure S5:**
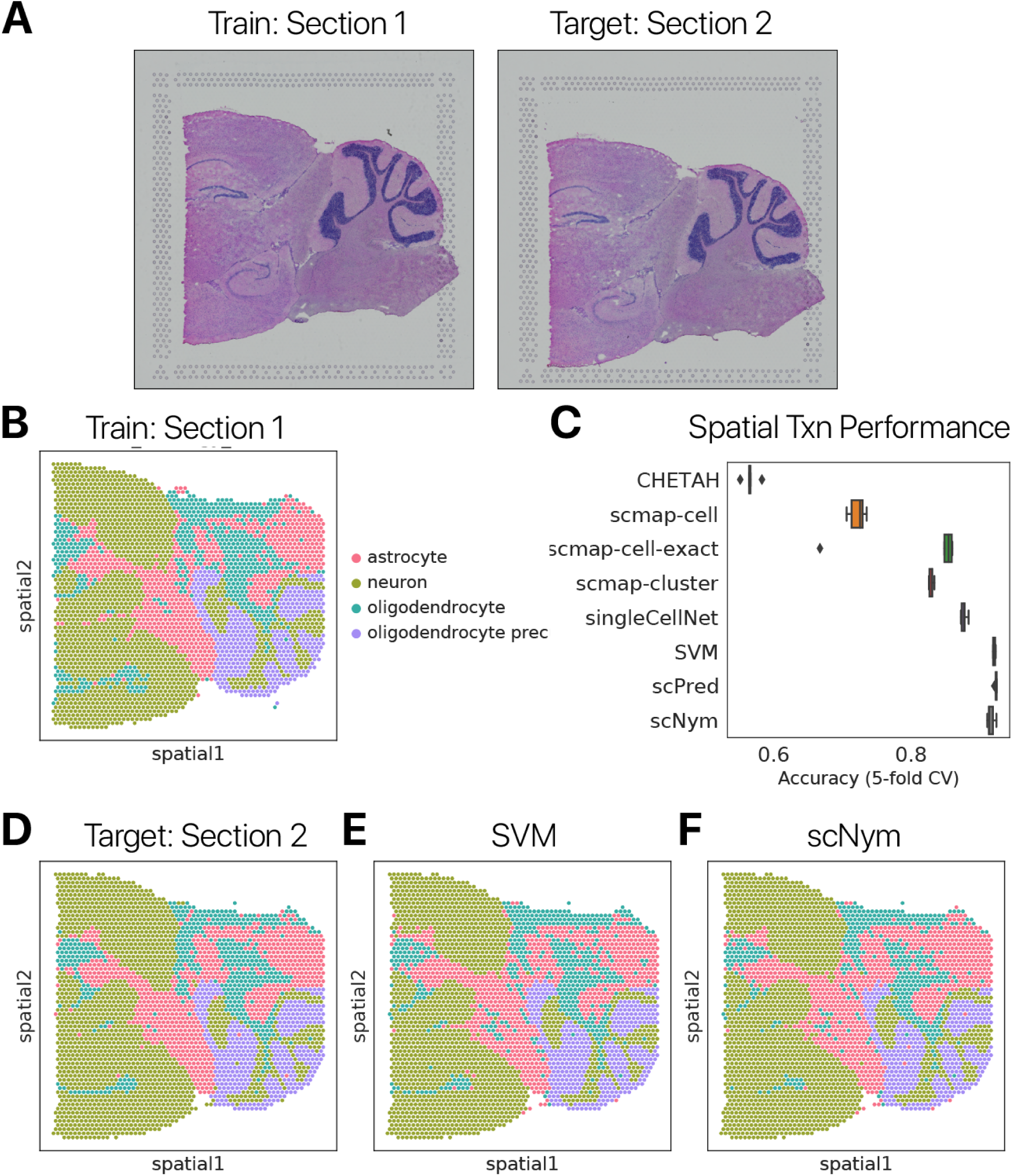
scNym transfers regional identity annotations across spatial transcriptomics data sets. **(A)** Images of training and target mouse brain sagittal-posterior sections profiled with spatial transcriptomics. **(B)** Ground truth regional identity annotations for the training section. **(C)** Accuracy score comparison of scNym to seven other state-of-the-art cell type classifiers. scNym models shown competitive performance relative to baseline methods, modestly less accurate than scPred and SVM approaches. **(D)** Ground truth regional identity annotations in the target section, alongside **(E)** SVM predictions and **(F)** scNym predictions.

**Figure S6:**
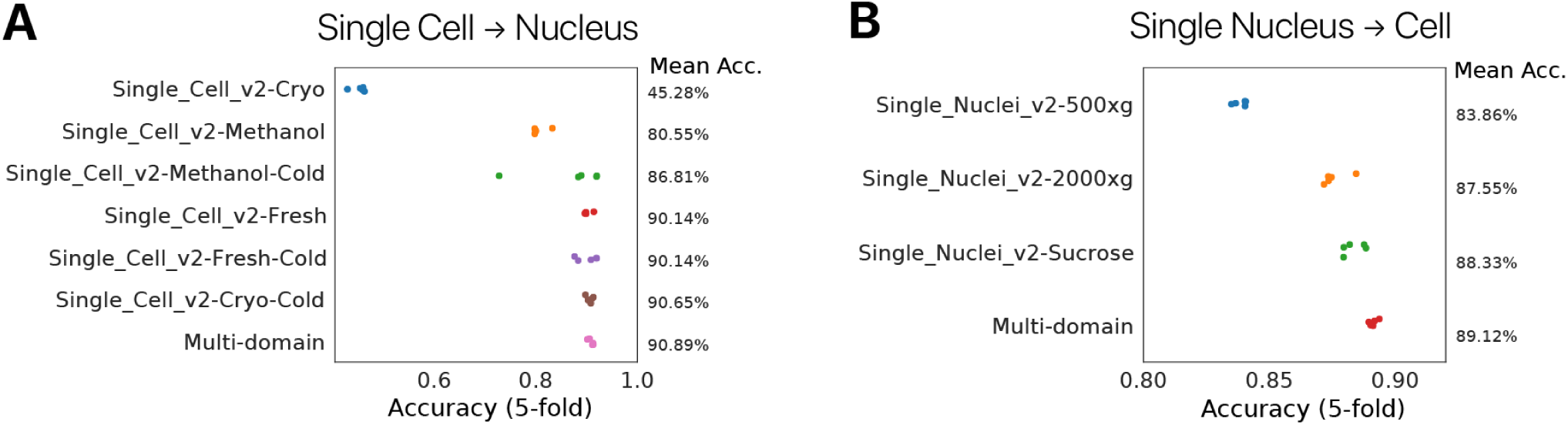
Multi-domain training improves performance of cross-technology annotation transfer in the mouse kidney. We trained scNym models using each training domain individually, or all domains through multi-domain training for both a single cell to single nucleus and single nucleus to single cell annotation transfer task. **(A)** Performance of different training approaches for the single cell to single nucleus task. Multi-domain models outperform models trained on any single domain. We cannot determine the best single training domain without access to ground truth target labels, so multi-domain training performs better than a realistic scenario where a user must select a single training domain *a priori*. **(B)** Similar results were obtained for single nucleus to single cell transfer tasks. Here, the benefit of multi-domain training is more dramatic than in the single cell to single nucleus case (Wilcoxon Rank Sum *p* < 0.01 for Multi-domain vs. Single Nuclei v2 - Sucrose).

**Figure S7:**
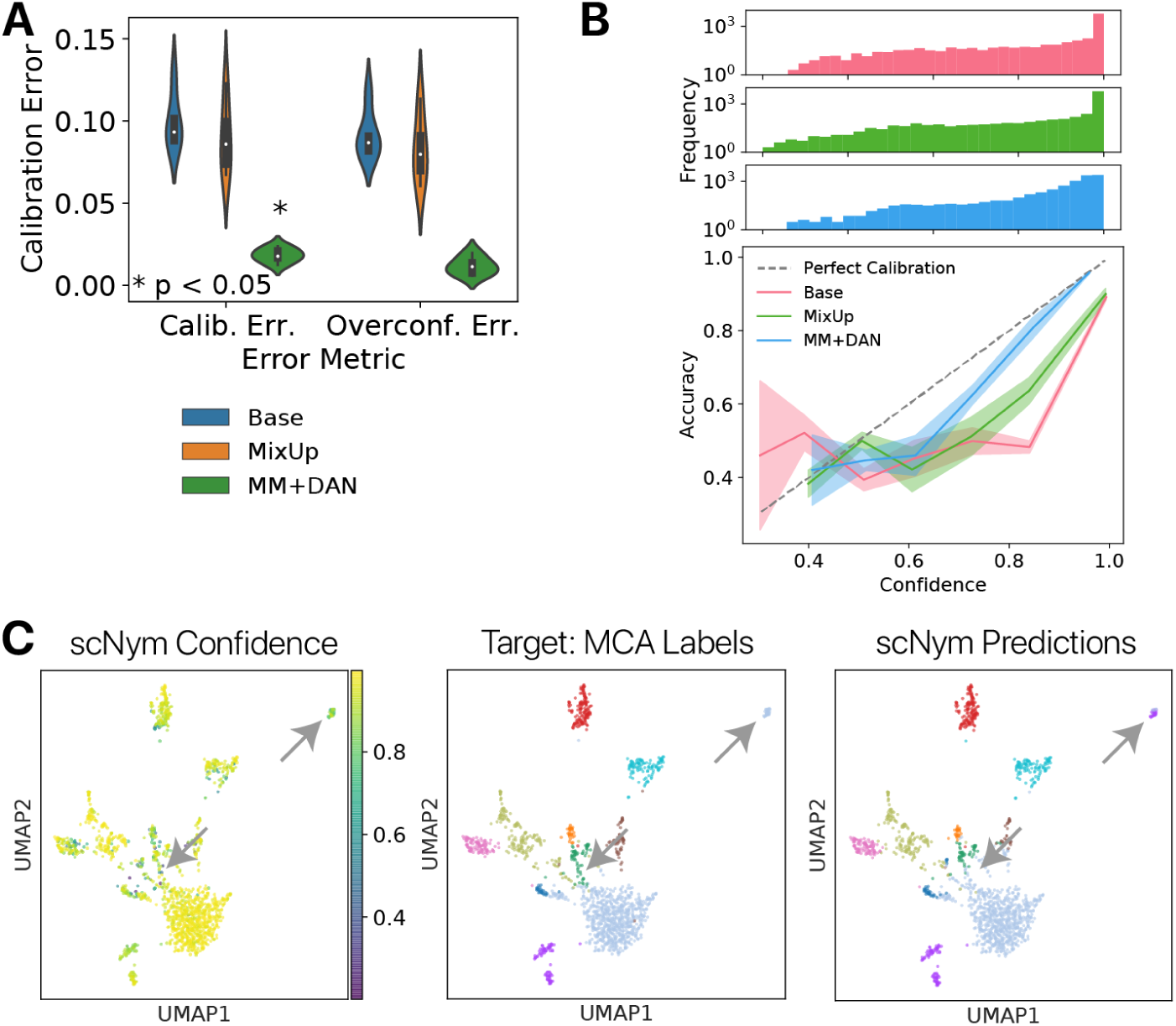
Semi-supervision improves the calibration of scNym models. **(A)** Expected calibration error (ECE) and overconfidence error (OE) scores for scNym models trained on the cross-stimulation benchmark without MixUp or MixMatch (Base), with MixUp on labeled data only (MixUp), or with the full MixMatch (MM) and domain adversarial network (DAN) procedure (MM + DAN). The MixMatch + DAN procedure significantly reduces the expected calibration error (*p* < 0.05, Wilcoxon Rank Sums). **(B)** Calibration curves for different scNym model configurations comparing the model confidence scores to observed classification accuracies. scNym models with MixUp, MixMatch, and DAN ablated (“Base”) show overconfident calibration at the high end. Confidence frequency histograms for each of the model configurations are shown above. **(C)** UMAP projection of model confidence scores, ground truth cell labels, and scNym predicted labels. We observed that incorrect predictions have lower confidence scores (arrows).

**Figure S8:**
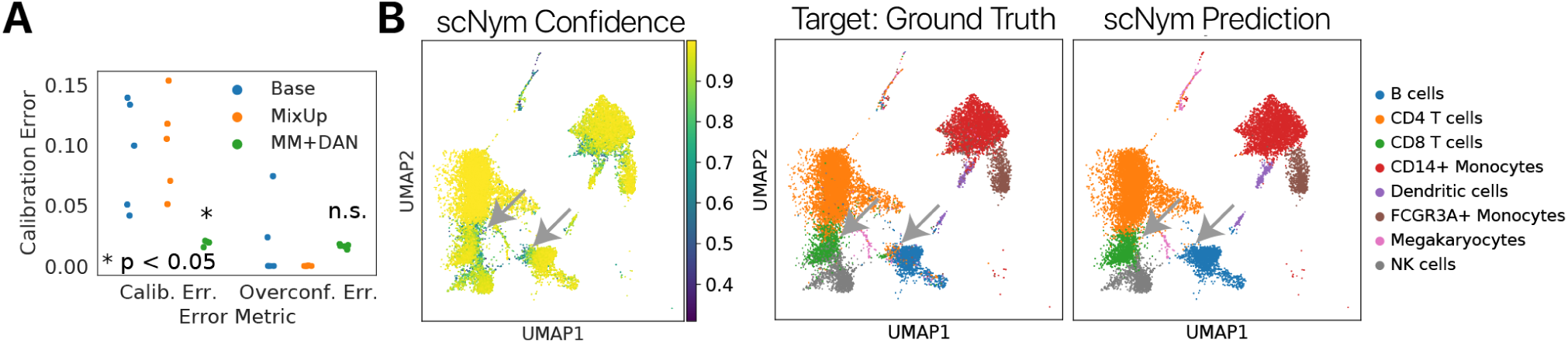
scNym model confidence highlights cells for manual curation. **(A)** Expected calibration error (ECE) and overconfidence error (OE) scores for scNym models trained on the cross-stimulation benchmark without MixUp or MixMatch (Base), with MixUp on labeled data only (MixUp), or with the full MixMatch and domain adversarial network (DAN) procedure (MM + DAN). The MixMatch + DAN procedure significantly reduces the expected calibration error (*p* < 0.05, Wilcoxon Rank Sums). **(B)** Confidence of the scNym model predictions projected in the UMAP embedding, alongside ground truth cell type annotations and scNym predictions. Low confidence scores identify incorrectly classified cells for manual curation (arrows).

**Figure S9:**
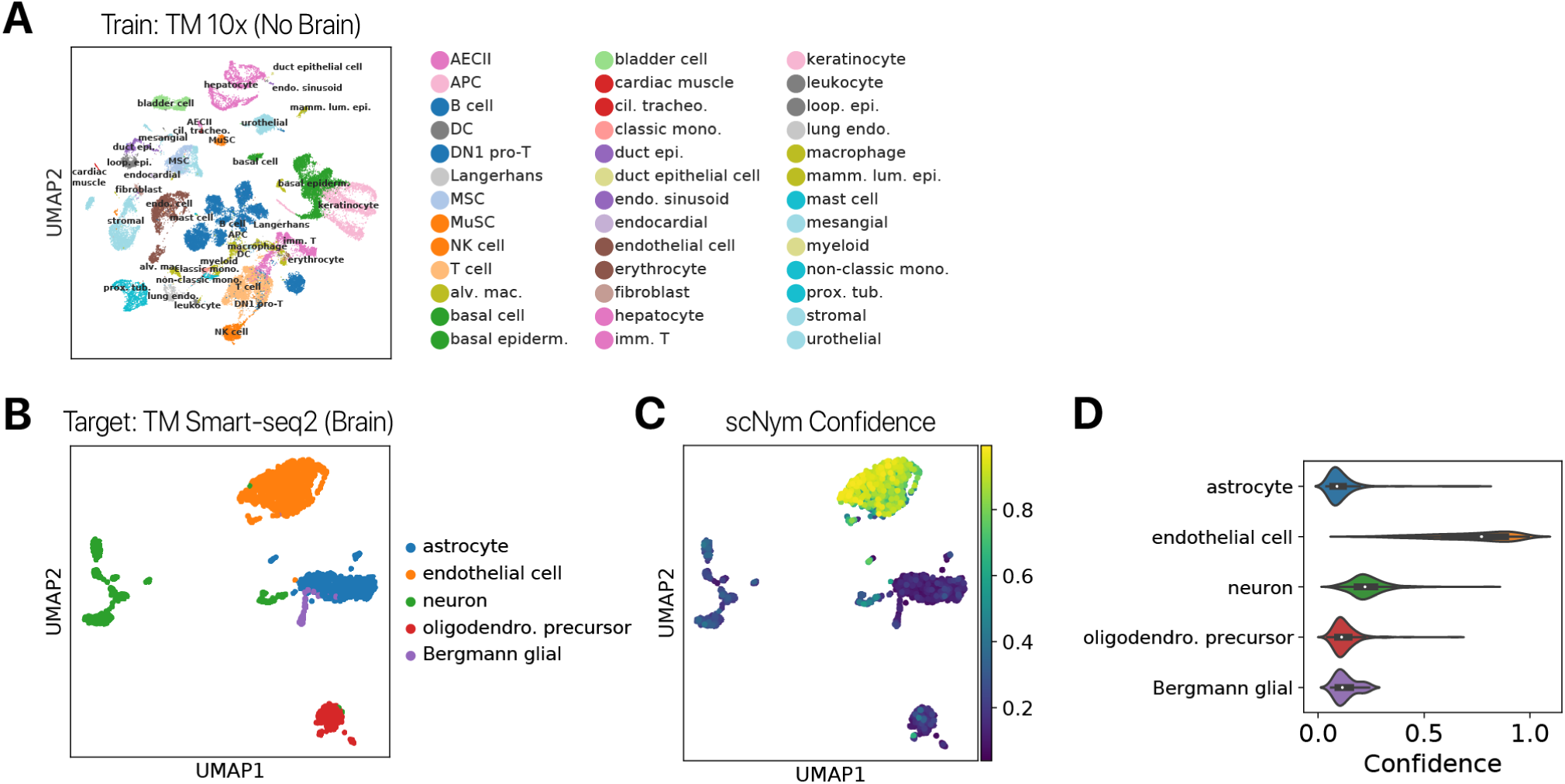
Confidence scores from pre-trained scNym models highlight new cell type discoveries. We trained an scNym model to transfer labels from the Tabula Muris 10x atlas to the Smart-Seq2 atlas, excluding brain tissue from the training and target sets. We subsequently predicted cell types for cells in the Smart-seq2 Brain (Non-Myeloid) tissue. **(A)** UMAP projections of ground truth cell type labels in the *Tabula Muris* 10x training data. Note that the training data does *not* contain any neurons, astrocytes, or other glia. **(B)** Ground truth cell type annotations in the Tabula Muris brain tissue captured using Smart-Seq2. Note that astrocyte, neuron, olidodendro. precursor, and Bergmann glia represent new, unseen cell types. **(C)** scNym model confidence scores projected in the Tabula Muris brain data. New cell types receive low confidence scores, while endothelial cells are correctly predicted with high confidence because they are present in the training data. **(D)** Distribution of confidence scores in **(C)** for each ground truth cell type.

**Figure S10:**
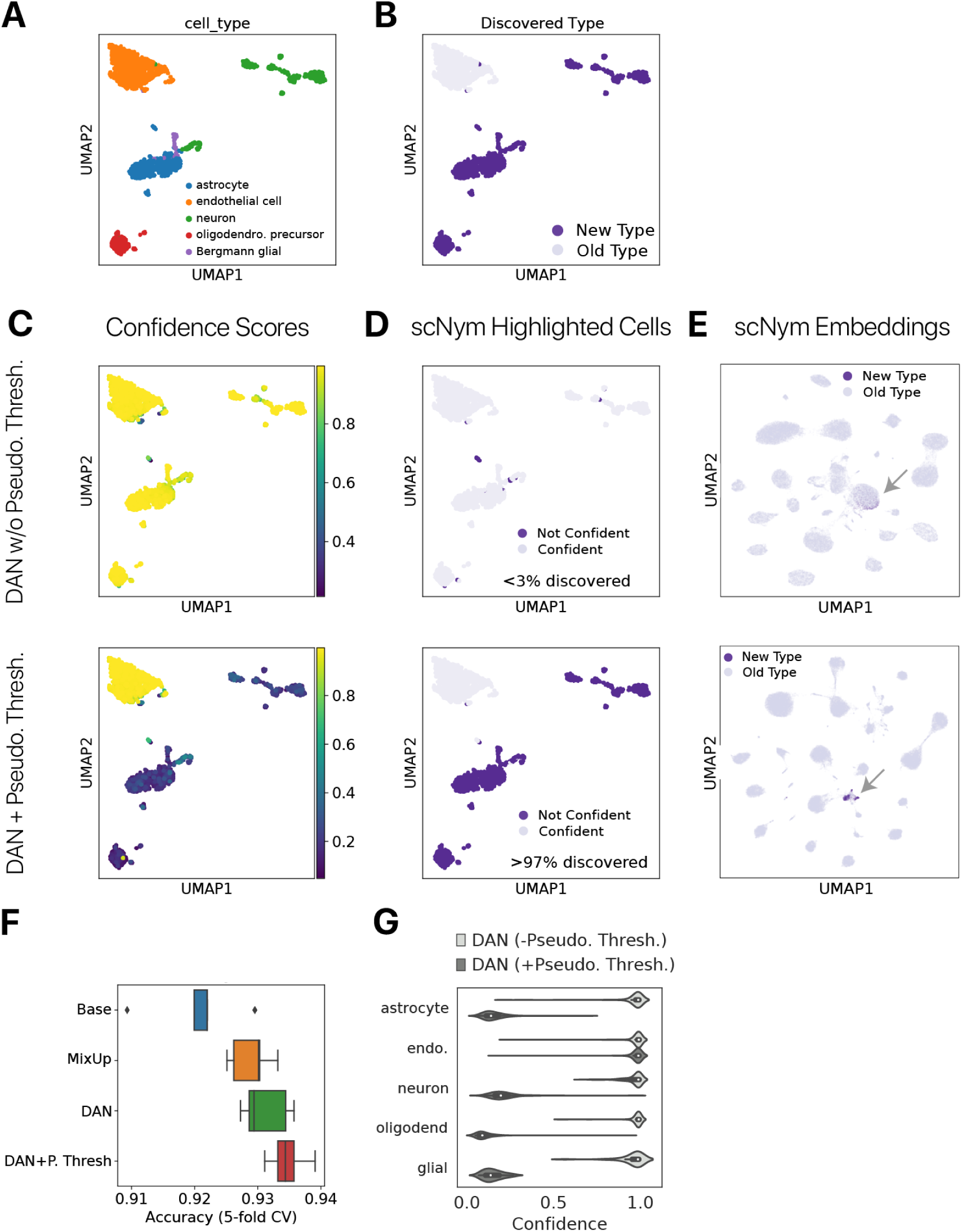
scNym confidence scores highlight new cell type discoveries after semi-supervised training. We trained a scNym models with and without pseudolabel thresholding to transfer labels from the Tabula Muris 10x atlas to the Smart-Seq2 atlas, *including* Brain tissue in the target set but not the training set. **(A)** Ground truth cell type annotations in the Tabula Muris brain tissue captured using Smart-Seq2. Note that astrocyte, neuron, olidodendro. precursor, and Bergmann glia represent new, unseen cell types. **(B)** New cell types “discovered” in this simulated experiment are highlighted in purple. **(C)** scNym confidence scores from models are displayed in the UMAP projection. **(D)** Cells highlighted as potential discoveries by thresholding scNym confidence scores (< 0.5). Models with pseudolabel thresholding correctly identify the majority (> 97%) of cells with novel types. **(E)** scNym embeddings. Pseudolabel thresholding allows new cell types to occupy a distinct region in the embedding (arrows). **(F)** Accuracy of cell type predictions for scNym models. Pseudolabel thresholding improves performance when new cell types are present. **(G)** Distribution of confidence scores for each ground truth cell type for scNym models. Models trained with pseudolabel thresholding correctly provide low confidence scores to new cell types, while models without pseudolabel thresholding incorrectly provide uniformly high confidence

**Figure S11:**
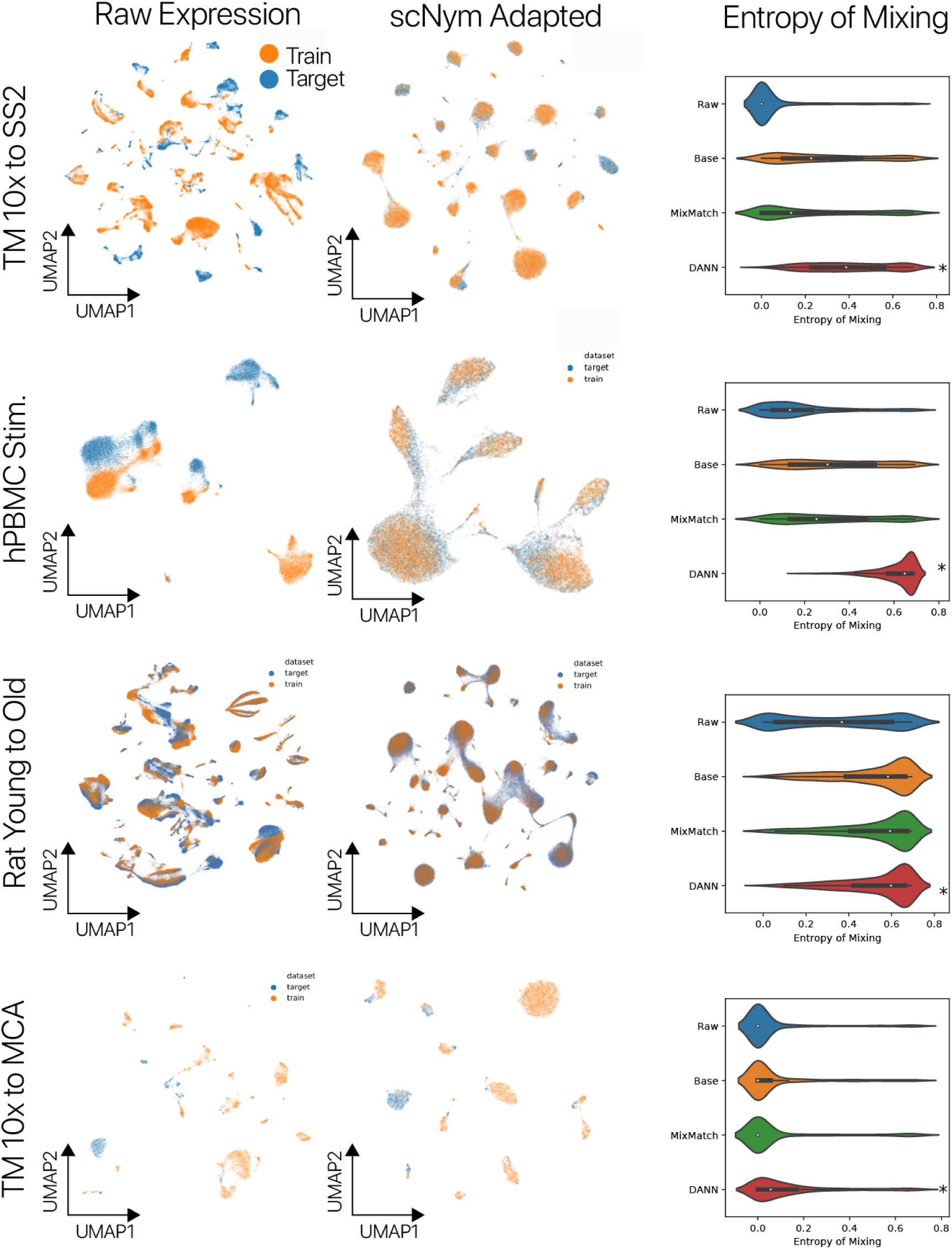
Adversarial scNym models adapt training and target domains. scNym models with MixMatch semi-supervision and a domain adversary were trained for each of our benchmark tasks. Raw gene expression embeddings derived using PCA and UMAP on normalized gene expression counts are shown on the left for each benchmark dataset. We have labeled the target and training domains with colors. In the center column, scNym model embeddings from the penultimate neural network layer are projected using PCA and UMAP. Qualitatively, we found that training and target datasets are more intermixed in the scNym embeddings. We confirmed this observation quantitatively by computing the entropy of batch mixing (right) on raw gene expression embeddings (Raw, left column embeddings), base scNym model embeddings (Base), scNym models with MixMatch but no domain adversary (MixMatch), and our complete scNym model with MixMatch and a domain adversary (DANN, center column embeddings). scNym models with MixMatch and a domain adversary were significantly more mixed than raw gene expression embeddings for each benchmark task (*p* < 0.05 Wilcoxon Rank Sums). This indicates that scNy2m5 models are effectively adapting across domains.

**Figure S12:**
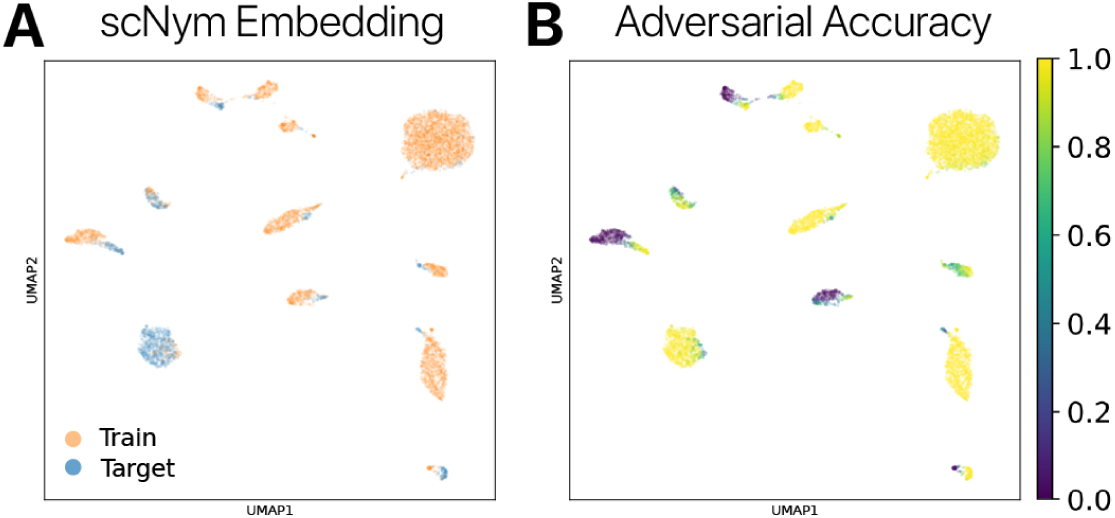
Domain adversarial training does not lead to aberrant mixture of training and target sets. We trained semi-supervised adversarial scNym models to transfer labels from the Tabula Muris 10x atlas to the Mouse Cell Atlas (Microwell-Seq). Despite the large class imbalance across the training and target domains, we found that domain adversarial training with a low weight (*λ*_adv_ = 0.1) does not lead to “overmixing” of the train and target domains in the embedding. **(A)** Train and target datasets are nearby in a series of cell type clusters, rather than collapsed into a single mode to beat the adversary without effective classification of cell types. **(B)** Adversary domain classification accuracy is presented as a color gradient in the embedding, smoothed over the nearest neighbor graph (*k* = 10). The adversarial accuracy is still high in many regions of the embedding after model convergence, confirming that the supervised loss is sufficient to counterbalance the gradients of the adversary to find high performance solutions.

**Figure S13:**
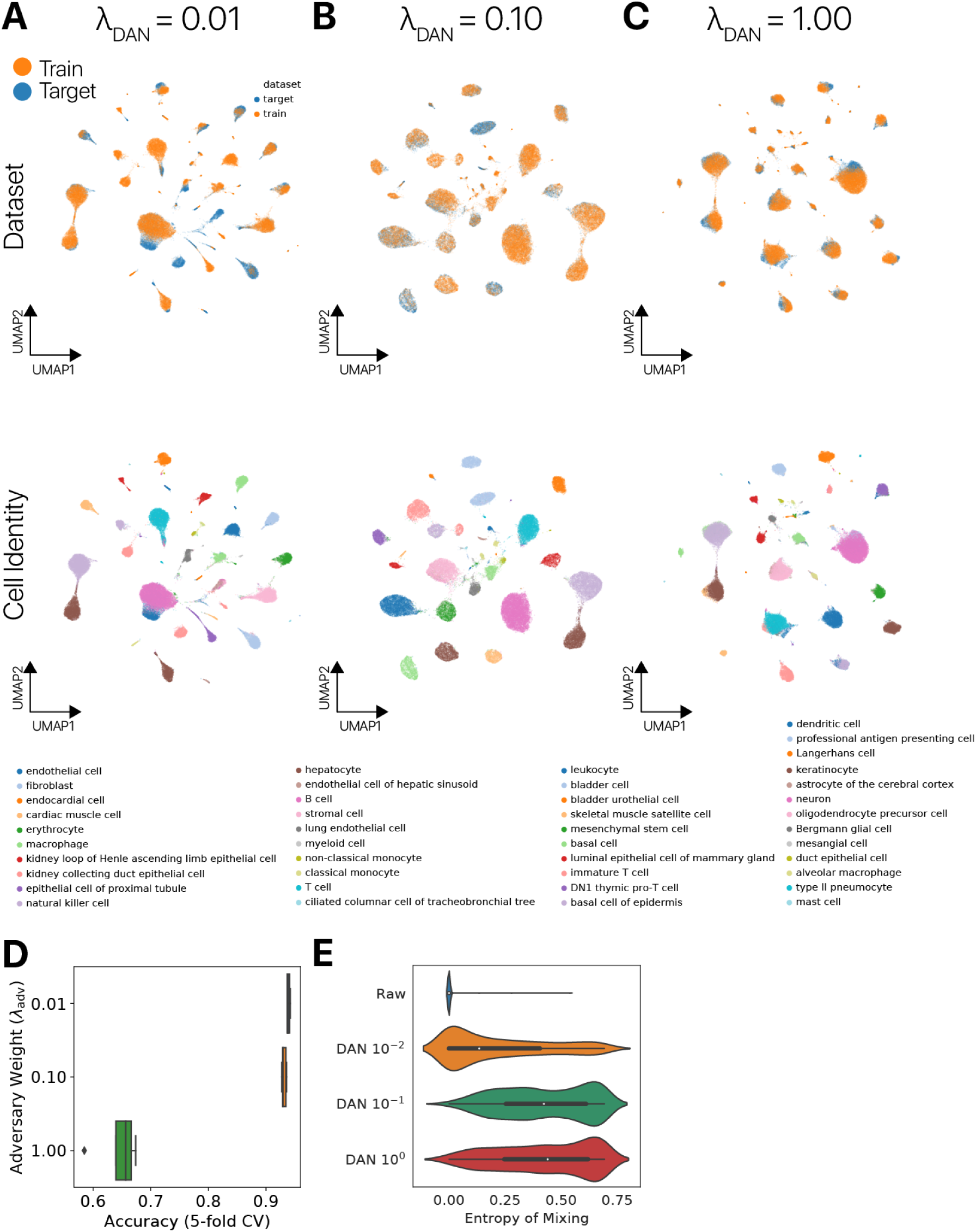
Exploration of domain adversary parameter space. We trained scNym models on the new cell type discovery experiment (Tabula Muris 10x without Brain to Smart-Seq2 with Brain) using a range of domain adversary loss weights. We compared the embeddings of these models after 80 epochs of training by visualizing with UMAP and labeling the dataset of origin and ground truth cell type for each observation. **(A, B)** Models trained with low domain adversary weights relative to the classification loss adapt across domains while allowing cell types to segregate in the latent space. **(C)** Models trained with high domain adversary weights relative to the classification loss collapse multiple cell types to a single mode in the latent space (note overlap of colors in Cell Identity plots). **(D)** Classification accuracy as a function of adversarial loss weight. Large adversarial weights reduce classification performance, likely because cell types are merged together in the embedding to compete with the adversary as in **(C). (E)** Entropy of mixing as a function of DAN weight. Raw indicates the raw gene expression embedding. Higher domain adversary weights increase the entropy of mixing in the embedding as predicted.

